# Intragenic suppressor screen of YHB identifies novel and known loss-of-function alleles of Arabidopsis phytochrome B

**DOI:** 10.64898/2026.05.07.723358

**Authors:** Wei Hu, Nathan C. Rockwell, J. Clark Lagarias

## Abstract

The red/far-red sensing photoreceptor phytochrome B (phyB) governs multifaceted plant development and responses to light and temperature stimuli. PhyB photoconversion between red-absorbing, inactive Pr and far red-absorbing, active Pfr states, imparted by its covalently bound bilin chromophore, enables rapid switching and plasticity of phyB signaling activities. The phyB^Y276H^ variant (YHB) is photochemically inert but adopts a constitutively active Pfr-like structure regardless of light conditions, which becomes a versatile model to dissect phyB signaling mechanisms. Here, we conducted a large-scale EMS mutagenesis screen on *YHB*-expressing transgenic lines, mining intragenic suppressor mutations that would unveil critical residues for phyB structure-function relationships. Comparative analyses of 26 nonsense variants suggested modular organization of phyB overall structure and dispensability of the C-terminal HKRD domain for phyB signaling. Amongst fourteen novel and nine known loss-of-function missense variants identified herein, G284E was of particular interest for its fully suppressed constitutive activity in darkness and its restored photochemistry and light responsiveness. The G284E mutation was further tested to also nullify another constitutively active phyB^Y303V^ allele by eliminating chromophore attachment. P309L was the sole variant identified which fully suppressed YHB in both dark and light conditions. C402Y profoundly elicited YHB protein instability. Three variants G118R, C402Y and G538D markedly reduced chromophorylation levels of YHB. Although the chromophore binding site variant C357Y was a strong loss-of-function allele, it retained residual signaling activity with respect to PIF3 protein turnover in dark-grown seedlings, presumably due to its ability to noncovalently bind chromophore. Two tandem prolines (P799, P800) proved critical to YHB structural integrity/stability as well as signaling activity. In summary, these diverse variants shed new insights into multiple levels by which the YHB (and thereby phyB) signaling is initiated, tuned, and disseminated.

## Introduction

Arabidopsis phytochrome B (phyB) is a red/far-red light photoreceptor that regulates plant growth and development in response to changing light conditions. Structurally, phyB functions as a homodimer, with each subunit comprising an N-terminal photosensory module—containing NTE, PAS, GAF, and PHY domains that house the covalently linked phytochromobilin (PØB) chromophore—and a C-terminal regulatory module involved in dimerization and signal transduction (Rockwell et al., 2006; Hughes and Winkler, 2024). Upon absorbing red light, phyB undergoes a reversible photoconversion from the inactive Pr form to the active Pfr form, triggering conformational changes that promote its accumulation in the nucleus. There, phyB interacts with transcription factors such as PIFs (phytochrome-interacting factors), leading to their inactivation and degradation to alter gene expression (Cheng et al., 2021). PhyB also senses temperature via its thermal Pfr-to-Pr reversion, which inactivates the signaling-active state (Jung et al., 2016; Legris et al., 2016) and the interaction with PIFs (Yi et al., 2024). Together, light and temperature signals are integrated by phyB to regulate many aspects of plant growth and development including seed germination, shade avoidance, and flowering time.

Substitution of a conserved tyrosine to histidine within the phytochrome GAF domain was first identified in the cyanobacterial phytochrome Cph1 through a directed evolution screen (Fischer and Lagarias, 2004). The resulting variant, Cph1^Y176H^, shows low photoactivity but exhibits strong red fluorescence. The same mutation, Y276H, in Arabidopsis phyB (herein referred to as YHB) also yields a red light–absorbing, Pr-like state that is strongly fluorescent (Fischer et al., 2005). Despite its Pr-like spectral properties, YHB behaves physiologically as though it is in the Pfr-like state, displaying constitutive activity *in planta* regardless of light conditions (Su and Lagarias, 2007; Hu et al., 2009). This behavior aligns with recent cryo-EM findings, which reveal that the overall structure of YHB closely resembles that of the active phyB-Pfr form, even though its bound PØB chromophore remains in a Pr-like ZZZssa configuration (Wang et al., 2024; Jia et al., 2025). Structurally, the H276 residue also mimics the positioning of Y276 in phyB-Pfr. In the Pr state (Li et al., 2022), such positioning would create a steric clash with the chromophore, forcing it to shift within the binding pocket. This displacement mirrors what is observed in phyB-Pfr and drives a conformational change in the PHY tongue from a beta-sheet to an alpha-helix, an established hallmark of the Pfr state in both plant and bacterial phytochromes (Yang et al., 2008; Anders et al., 2013; Takala et al., 2014; Wang et al., 2024). Because YHB remains constitutively active irrespective of light, it has become a valuable tool for studying phyB signaling pathways (Galvao et al., 2012; Jones et al., 2015; Jung et al., 2016; Huang et al., 2019; Chen et al., 2022) and has been widely used in efforts to engineer new plant traits (Hu and Lagarias, 2017; Alves et al., 2020; Hu et al., 2020; Battle et al., 2024).

As a central regulator of plant responses to both light and temperature signals, phyB has been extensively targeted through multiple approaches to identify alleles that either weaken or enhance its function. Early genetic screens uncovered strong loss-of-function (LOF) mutations at the endogenous *PHYB* locus (Reed et al., 1993; Bradley et al., 1996; Krall and Reed, 2000). Subsequent chemical mutagenesis screens of PHYB-overexpressing transgenic lines revealed additional LOF alleles, including both missense and nonsense mutations (Wagner and Quail, 1995; Chen et al., 2003; Oka et al., 2008). A yeast reverse-hybrid mutagenesis screen further identified variants that disrupt the interaction between phyB and PIF3 (Kikis et al., 2009). Beyond these approaches, both LOF and gain-of-function (GOF) alleles have been generated through structure-guided mutagenesis (Zhang et al., 2013; Wang et al., 2024), in vitro photochemical analyses (Su and Lagarias, 2007; Burgie et al., 2014; Jeong et al., 2016), targeted mutation of residues involved in posttranslational modifications (Medzihradszky et al., 2013; Nito et al., 2013; Sadanandom et al., 2015; Viczian et al., 2020; Zhao et al., 2023), and mining natural variation across different accessions (Maloof et al., 2001; Filiault et al., 2008). The growing catalog of phyB mutations (Bae and Choi, 2008; Klose et al., 2020; Chen et al., 2024) has provided key insights into diverse aspects of phyB biology, including chromophore assembly, photoconversion, thermal reversion, protein stability, nuclear localization, photobody formation, regulation by posttranslational modifications, and interactions with downstream signaling partners.

In this study, we leveraged new plant lines expressing the dominant, constitutively active YHB allele to identify intragenic suppressor variants via a large-scale genetic screen. Since YHB does not exhibit Pr-to-Pfr photoconversion, second site missense mutations accelerating thermal reversion and/or destabilizing the Pfr state were not expected to be identified. Hence, this screen should enrich for mutations affecting chromophorylation levels, protein stability, structural configuration, interaction with signaling partners, and subcellular migration. Our screen has identified both missense and nonsense alleles, shedding new insight into the structural basis of phyB signaling. Remarkably, the G284E variant was found to specifically restore photochemistry and thereby deactivate the constitutive activity of YHB. The Glycine-284 residue was also critical in underpinning another constitutively active allele phyB^Y303V^ (YVB) (Jeong et al., 2016).

## Results

### EMS mutagenesis of single-insertion *YHB* transgenic lines

Our screen required the isolation of new single-insertion *YHB* lines in the *phyB-5* null mutant background, because a prior elite *YHB*^*g*^*/phyA-201phyB-5* line #5 (Su and Lagarias, 2007) had tandem insertion of the YHB^g^ transgene (Hu et al., 2009) which prevented identification of intragenic mutations. We therefore introduced a synthetic eYHB-3xFLAG coding sequence (Hu and Lagarias, 2017) driven by the constitutive CaMV 35S promoter into the *phyB-5* mutant background. Through genetic analyses and mapping of T-DNA insertion loci, we obtained three single-insertion *35S::eYHB-3xFLAG*/*phyB-5* lines (#3, #4, #11) - all of which exhibited stable GOF constitutively photomorphogenesis (cop) phenotypes (Fig. S1). To offset potential bias of transgene loci on mutagenesis efficiency, we employed two lines for the screen - line #3 (T-DNA inserted near the telomere region of Chr 4) and line #4 (T-DNA inserted near the center region of Chr 1). Approximately 50,000 seeds in total from both lines were chemically mutagenized, and T2 seedlings were screened for the loss of the GOF short hypocotyl phenotype using the protocol depicted in Fig. S2. Our screen rejected extragenic suppressor mutants in the chromophore biosynthesis genes HY1 and HY2. These and other extragenic suppressors were not the focus of this study. Intragenic suppressor mutants isolated from line #3 were named as NA# series and those from line #4 as NB# series. A total of 26 nonsense and 23 missense mutations were identified as summarized in Table S1.

### Phenotypic characterization of nonsense suppressor variants reveals structural modularity of phyB

As shown in Fig. 1A and Table 1, the 26 nonsense mutations were distributed throughout the YHB coding sequence, with most mutations falling within the GAF and PHY domains and none in the NTE domain. Two widely studied LOF *phyB-1* (Q448*) and *phyB-9* (W397*) null alleles were among these nonsense mutations. Given the exclusive G/C-to-A/T mutation pattern elicited by EMS, only the codons of Glutamine (Q), Tryptophan (W) and Arginine (R) could be mutated to stop codons. Of the 81 potential target sites in YHB, 26 nonsense mutations were identified - eight of which were independently recovered multiple times (mutation events *n* = 41) (Figs. 1A, S3).

**Fig. 1.**
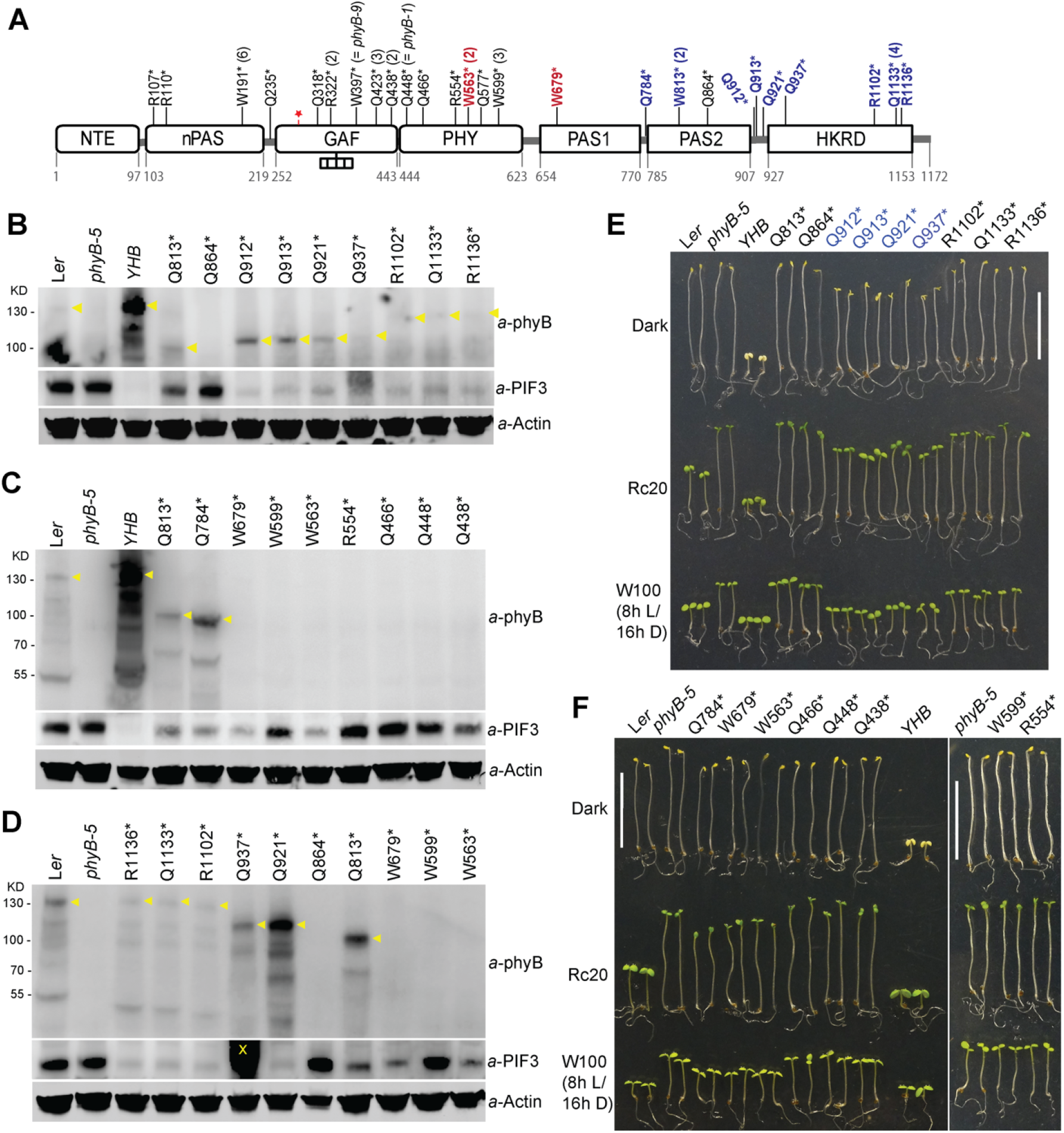
YHB nonsense suppressor mutations. (**A**) Schematic representation of PHYB domain structure and identified nonsense mutations. The red star denotes the Y276H mutation, the tandem rectangle denotes the attached bilin chromophore, and numbers in parentheses indicate the number of independent isolation of the same mutant. Variant lines with both reduced PIF3 protein levels and accumulation of truncated YHB are indicated in bold blue, while those with reduced PIF3 protein levels and undetectable truncated YHB in bold red. (**B-D**) Immunoblot assays of truncated YHB variant and PIF3 protein levels from dark-grown seedlings for selected nonsense suppressors (C-terminal to Q438*). Yellow arrowheads depict positions of phyB/YHB and truncated YHB variants. In (D), the parental YHB line was excluded to improve signal sensitivity from three weakly expressed nonsense mutants R1136*, Q1133* and R1102*. “X” mark indicates a strong non-specific band associated with Q937* that overly masks the weak PIF3 band below, which is better seen in panel (B). (**E-F**) Four-day-old seedlings grown in the dark, under red light (20 µmol m^-2^ s^-1^, Rc20) and under short-day white light (W100) conditions, bars = 1 cm. Q912* ∼ Q937* mutants labeled in blue exhibit discernable cop phenotypes in darkness and have the strongest light growth phenotypes among all nonsense mutants.

Mutational effects of these nonsense variants were holistically evaluated by plant phenotyping and by immunoblotting of seedling extracts to measure levels of truncated YHB and PIF3 proteins. A monoclonal antibody B1, targeted to an epitope within the NTE domain of PHYB (Wagner et al., 1996), was used to detect accumulation of any truncated YHB variant in dark-grown seedling extracts. The same extracts were probed with antibodies to PIF3 (Fig. 1B-D). Since PIF3 is a well-known target for phyB-dependent turnover, its level served as a sensitive molecular indicator of phyB activity (Al-Sady et al., 2006; Hu and Lagarias, 2017, 2024).

Immunoblotting revealed a broad range of residual expression of truncated YHB proteins - all of which accumulated significantly lower levels than that present in the *YHB* parental line (Fig. 1B-D). This shows that the synthesis and/or stability of the truncated proteins were reduced in all lines. Correspondingly, PIF3 was significantly degraded in the variants expressing detectable truncated YHB. Notably, a cluster of three variants (Q912*, Q913* and Q921*) truncated between the PAS2 and HKRD domains, and one variant (Q937*) truncated at the far N-terminal region of the HKRD domain, expressed truncated YHB at levels higher than or similar to endogenous phyB in L*er* WT (Fig. 1B, 1D). These four nonsense alleles exhibited weak cop phenotypes in darkness and retained the ability to complement *phyB-5* mutant phenotypes of light-grown seedling and adult plants (Fig. 1E, S4). These results corroborate earlier studies which show that the HKRD domain is dispensable for phyB signaling (Krall and Reed, 2000) as long as the truncated protein is reasonably stable. Indeed, three other nonsense variants (R1102*, Q1133* and R1136*) which lack a small C-terminal portion of the HKRD domain proved to be unstable (Fig. 1B, 1D) and poorly complemented *phyB-5* morphologically (Fig. 1E, S4). That said, R1102*, Q1133* and R1136* did induce PIF3 protein turnover as effectively as Q912* ∼ Q937*, documenting their residual activity.

Among the 11 C-terminal nonsense mutations, Q864* was a phenotypically null allele because no truncated YHB was detectable nor was PIF3 turnover observed (Fig. 1B, 1D, 1E). Q864* plants also were phenotypically indistinguishable from the *phyB-5* mutant (Fig. 1E). Q864*’s truncation within the PAS2 domain is likely responsible for generation of an unstable protein. By comparison, both W813* (within the PAS2 domain) and Q784* (immediately N-terminal to the PAS2 domain) accumulated significant levels of truncated YHBs. These two variants, as well as W679* (within the PAS1 domain) and W563* (within the PHY domain) all induced significant PIF3 turnover (Fig. 1C, 1D). Because both W679* and W563* promoted PIF3 protein turnover and conferred discernable photomorphogenic growth phenotypes, some truncated YHB proteins must have been produced despite our inability to detect them immunochemically (Fig. 1C, 1D, 1F).

Five additional nonsense variants in the N-terminal PHY and GAF domains, W599*, Q577*, R554*, Q466*, Q448* (a.k.a. *phyB-1*) and Q438*, were tested to be authentically null both at the molecular and phenotypic levels (Fig. 1). The remaining eight variants further N-terminal to Q438* were phenotypically null as expected consistent with their incomplete chromophore-binding GAF domain and were classified as null alleles without further analysis.

### Phenotypic characterization of missense suppressor variants

As shown in Fig. 2A and Table 1, we identified nine previously known and 14 novel missense suppressor variants, some of which were recovered multiple times. No missense mutation was found within the NTE or HKRD domains. Novel suppressor mutations included G118E, P145L, D307N, C357Y, C402Y, G538D, P581L, E655K, A663V, A669V, G678E, P799L, P800L and G899E. Immunoblotting showed minor changes in the levels of most suppressor variants relative to YHB in the non-mutagenized parent line, with the notable exceptions of the R110Q, G118R and C402Y variants, which were greatly reduced (Fig. 2B, upper panel). After genetic backcrossing to clean up mutational backgrounds, however, the R110Q protein levels were comparable to the YHB protein level whereas the C402Y protein levels remained strongly reduced (Fig. S5). Immunoblotting indicated that the C402Y protein level was at least 8-fold less than the parental YHB level, which was still comparable to endogenous phyB levels in L*er* WT (Fig. S5A, B, C). By contrast, the *YHB*^*C402Y*^ transcript level was not reduced and indeed was even higher than that of the parental eYHB transcript (Fig. S5D). These results show that the C402Y protein product was unstable. Quantitative RT-PCR also revealed that R110Q, C357Y, C402Y and G767R mutations all mitigated YHB-dependent induction of the *STH* transcript and suppression of the *PORA* transcript in the dark-grown seedlings - two known target genes of YHB (Fig. S5D).

**Fig. 2.**
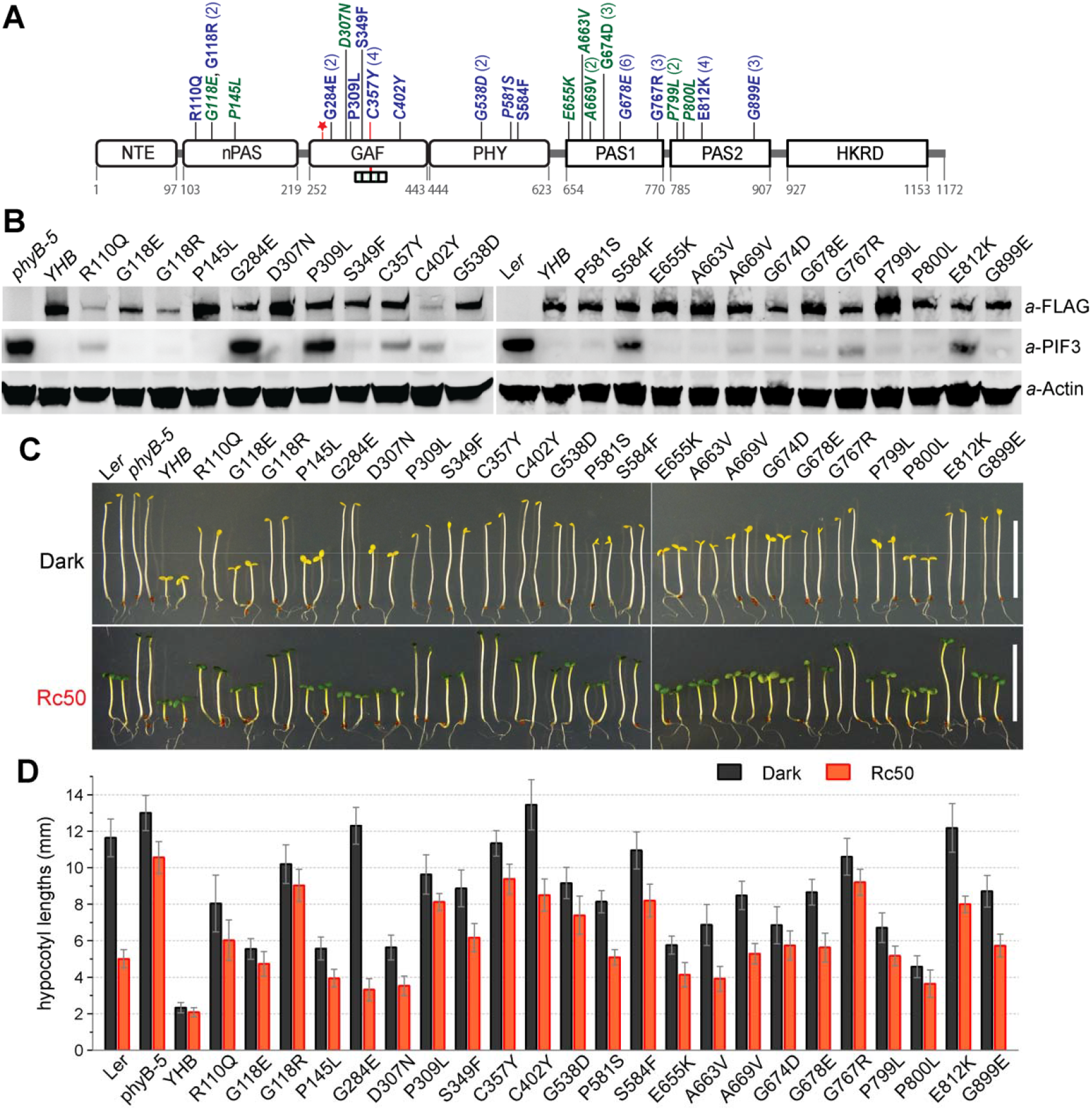
YHB missense suppressor mutations. (**A**) Schematic representation of PHYB domain structure and identified missense mutations. The red star denotes the Y276H mutation, the tandem rectangle denotes the attached bilin chromophore, and numbers in parentheses indicate the number of independent isolation of the same mutant. Mutations in blue confer strong suppression of YHB function in dark-grown seedlings, whereas mutations in green are weak suppressors (moderate cop phenotypes, with well-expanded cotyledons). Italicized mutations are novel ones identified in the current study. (**B**) Immunoblot assays of YHB-3xFLAG and PIF3 protein levels of dark-grown seedlings. (**C**) Four-day-old seedlings grown in the dark or under continuous red light (Rc50), bars = 1 cm. (**D**) Quantitative measurement of hypocotyl lengths of seedlings shown in (C), **n** = 10 ∼25.

Companion blots were also probed with the PIF3 antibody to evaluate the functional strength of YHB variants in dark-grown seedlings (Fig. 2B, middle panel). Based on the stability of PIF3, G284E and P309L were the two fully inactive variants. Other strong suppressors included four known LOF variants R110Q, S584F, G767R and E812K, as well as two novel variants C357Y and C402Y - all of which retained measurable, albeit reduced, PIF3 protein levels compared with dark-grown WT or *phyB-5*. Consistently, these eight strong suppressor lines were etiolated or nearly etiolated in the dark, revealing strong suppression of YHB function (Fig. 2C, 2D). Four other moderate ∼ strong suppressor variants, S349F, G538D, G678E and G899E, also profoundly promoted hypocotyl elongation and inhibited cotyledon unfolding/expansion of dark-grown seedlings, despite the strong reduction of PIF3 levels. The remaining variants (G118E, P145L, D307N, P581S, E655K, A663V, A669V, G674D, P799L and P800L) exhibited moderate *cop* phenotypes, showing opened cotyledons while retaining longer hypocotyls than the parent *YHB* line in darkness.

When grown under continuous red light (50 µmol m^-2^ s^-1^), all missense variant seedlings except G284E displayed LOF phenotypes consistent with their effects in the dark: strong suppressors had longer hypocotyls while weak suppressors had shorter hypocotyls (Fig. 2C, 2D). The G284E variant seedlings were etiolated in darkness indicating full suppression of the constitutive activity of YHB. They were surprisingly strongly responsive to red light. Thus, the LOF G284E mutation is conditional, with red light effecting reactivation of YHB signaling. As such, this mutant line is phenotypically similar to WT - inactive in darkness and activated by light.

Adult plant phenotypes of missense variants grown in short-day photoperiods were also examined for the impairment of YHB signaling. These analyses revealed P309L and C357Y to be the only two variants fully suppressing YHB activity, with both exhibiting elongated petioles and poorly expanded leaves similar to those of the *phyB-5* mutant (Fig. S6). Other variants displayed adult plant morphology in the range between L*er* WT and the *YHB* parent.

### Chromophore binding and photochemical properties of N-terminal missense suppressor variants expressed in *E. coli*

The thirteen missense alleles found in the YHB photosensory region were subcloned by PCR amplification, expressed as truncated N651 proteins in *E. coli* along with a PΦB-producing biosynthetic operon, and affinity purified. Whereas all of the variants yielded recombinant proteins of the expected molecular mass, pronounced heterogeneity was seen in many of them after SDS-PAGE. Notably, the G118E, G118R, C402Y and G538D variants exhibited two distinctly separated isoforms, while multiple species were also seen for the D307N, P309L, S349F and C357Y variants (Fig. 3A, bottom panel). Companion zinc blot analyses revealed that the slowest migrating species represented chromophore-bound holoprotein for those multiple-isoform variants except for C357Y (Fig. 3A, top three panels). Indeed, poor chromophorylation of the C357Y variant was expected, since Cys357 is the conserved site of chromophore binding in phyB. In addition, covalent PΦB binding was greatly reduced for G118E, G118R, D307N, P309L, C402Y and G538D variants in comparison to WT PHYB or YHB controls.

**Fig. 3.**
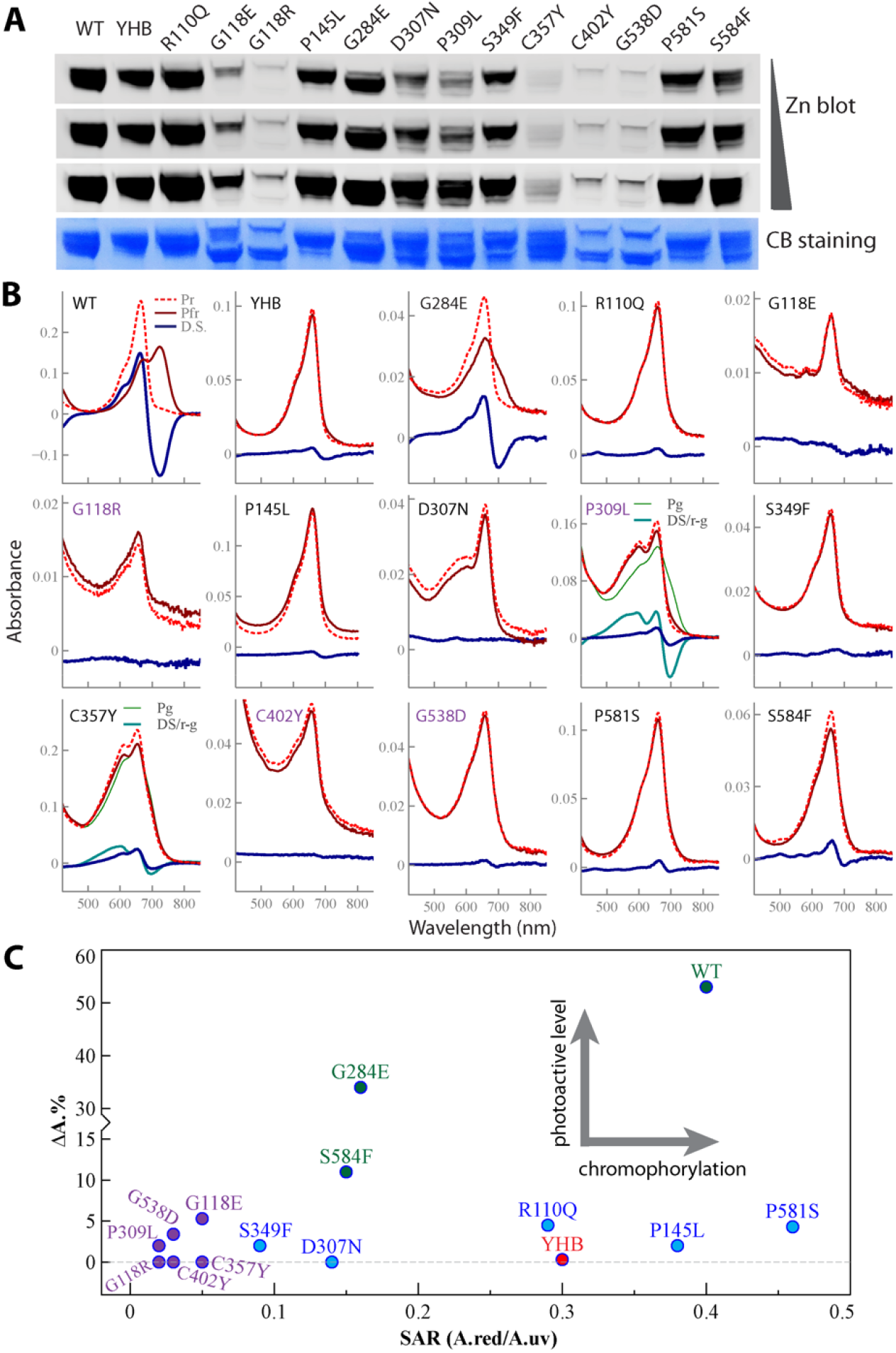
Zinc blot assay and photochemical properties of N-terminal missense variants expressed as truncated photosensory domain proteins. (**A**) Zinc blot assay for detecting covalently bound chromophore of recombinant proteins isolated from PØB-producing E. coli cultures. Exposure times of zinc blot was varied to better view low signal proteins; Coomassie Blue staining shows protein loading levels. (**B**) Spectral analysis of N651 recombinant proteins. Variant proteins in purple color were concentrated at least 10-fold to increase signal strength. P307L and C357Y had a secondary absorbance peak around 600 nm, so additional Pr-Pg(green) difference spectra were also presented. D.S. = Pr-Pfr difference spectra. (**C**) Normalized chromophorylation level vs photoactive levels of N651 recombinant proteins; SAR, specific absorbance ratio, = (Abs. of red peak)/(Abs. of UV peak) of Pr sample; ΔA.% = (Abs. of Pr - Abs. Pfr)/(Abs. of Pr) x 100% to depict photoconvertibility or Pr depletion level.

Examination of the absorption spectra of the 13 missense variants corroborated the results of zinc blot assays, with the notable exception of C357Y which yielded a strong red-absorbing product in the native state despite its poor covalent chromophore binding (Fig. 3B). Some variant proteins had to be concentrated to obtain reliable absorption data, because of their poor chromophorylation levels as determined by Specific Absorption Ratio measurements (SAR, Fig. 3C). As expected, most variants yielded red-absorbing holoproteins that were poorly photoactive like the YHB parent (Fig. 3B). Surprisingly, G284E was photoactive and exhibited a photochemical difference spectrum similar to WT phyB (Fig. 3B). This photochemical property is consistent with the plant phenotypes of this mutant line which exhibits no signaling activity in darkness and is responsive to red light (see above). Interestingly, the G284E mutation was previously recovered as a LOF allele of phyB (the phyBN651-GFP-GUS-NLS transgene) (Oka et al., 2008). Hence, the effect of the G284E mutation is context dependent, restoring light signaling activity in the YHB background while inhibiting light signaling in the WT phyB background. The S584F mutation also exhibited an increase in photochemistry yet failed to produce a signaling active Pfr form (Fig 3B, 3C), which is consistent with its LOF activity.

The D307N, P309L and C357Y variants were of interest because they all had a secondary absorption maximum in the yellow-orange region in the dark state, perhaps indicating the presence of deprotonated bilin species (Fig. 3B). Whereas D307N was photoinactive, both P309L and C357Y comprised mixtures of species with red/green and red/far-red photocycles. Their unusual photochemical behavior belied their inability to rescue light-dependent signaling (Fig. 2, S6). Indeed, P309L was one of the strongest YHB suppressors both in darkness and under all light conditions tested. C357Y was noteworthy because its non-covalent binding of chromophore conferred novel photochemical behavior (Fig. 3B) but did not restore light-dependent signaling activity (Fig. 2, S6).

### Missense variants abolish red fluorescence or alter subnuclear distribution of YHB as revealed by fluorescence confocal microscopy

Arabidopsis YHB is strongly fluorescent and forms a few large nuclear photobodies independent of light conditions (Su and Lagarias, 2007; Chen et al., 2022). We therefore used confocal microscopy to assess the subcellular localization of each missense variant - where detectable - in dark grown seedlings (Fig. 4; Fig. S8). As expected, photosensory domain variants that bound PØB poorly *in vitro* or those with restored photochemical activity were difficult to detect by fluorescence. These included the three strongest suppressor variants G284E, P309L and C357Y lacking detectable nuclear fluorescence, as well as G118R, C402Y, G538D and S584F exhibiting diffuse, barely detectable nuclear fluorescence. Apparent exceptions to this pattern included G118E and S349F, for which both small and large nuclear photobodies were detected, and D307N, where small photobodies or steady diffuse fluorescence were seen (Fig. 4). The other three photosensory domain variants R110Q, R145L and P581S that exhibited robust SAR values as recombinant proteins (Fig. 3C) could all be detected in the nucleus. P145L had both small and large photobodies, R110Q exhibited steady diffuse fluorescence with occasional small photobodies, and P581S only had weakly diffuse fluorescence.

**Fig. 4.**
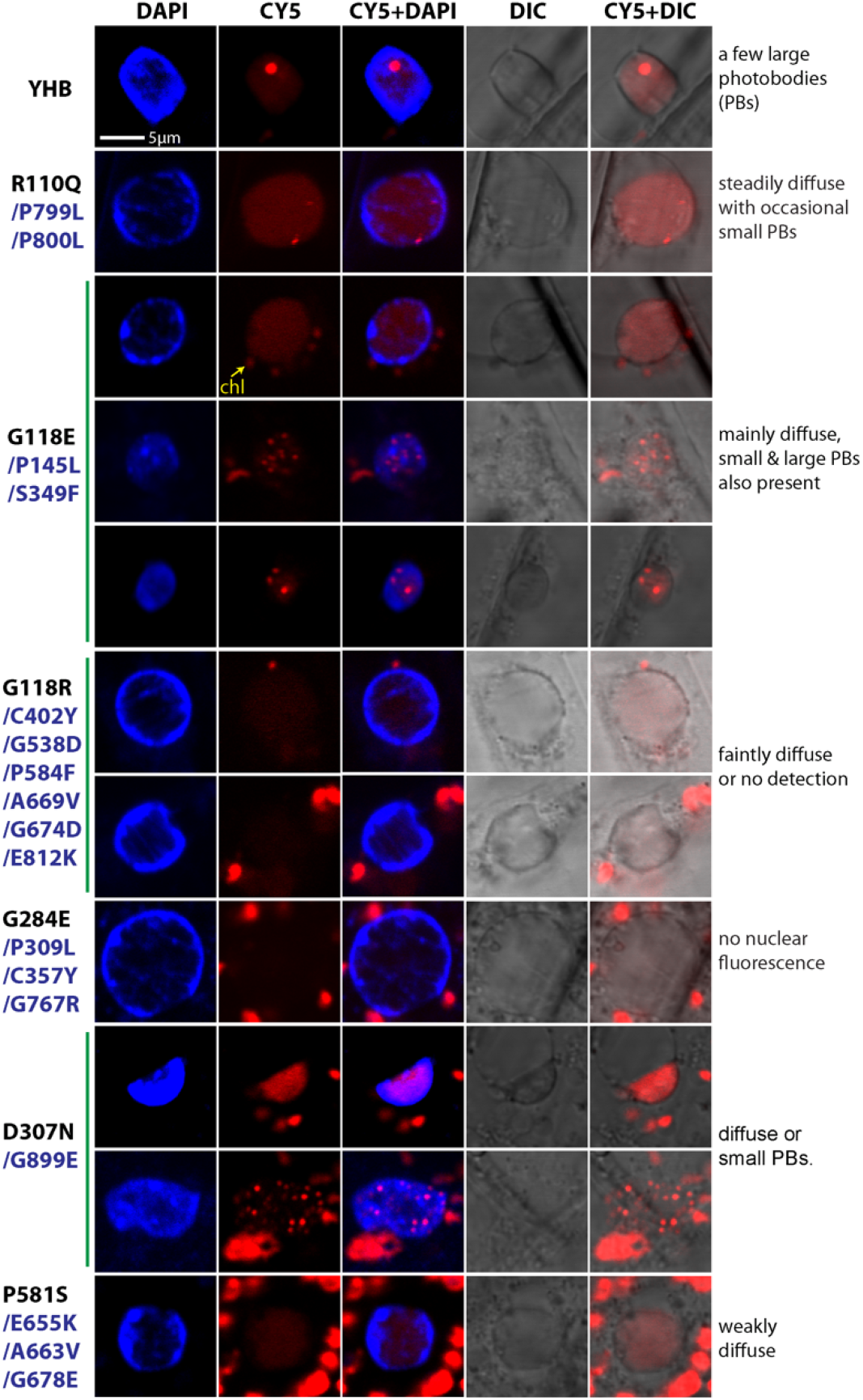
Missense mutations abolish red fluorescence of YHB or alter its subnuclear distribution pattern in dark-grown seedlings. The CY5 channel of confocal microscopy simultaneously captures red fluorescence emitted by YHB and chlorophyll. The nucleus is revealed by DAPI staining. Similar subnuclear distribution patterns are classified and represented by one missense mutation, and images for other mutations of the same pattern group are provided in Fig. S8. Varied patterns, if any, of individual classification group are shown.

Notable amongst the regulatory domain variants, G767R lacked any nuclear fluorescence (Fig. 4). Variants A669V, G674D and E812K exhibited nearly undetectable diffuse nuclear fluorescence, whereas E655K, A663V and G678E exhibited weakly diffuse nuclear fluorescence. By contrast, variants P799L and P800L had reliably strong signals, with diffuse nuclear fluorescence and occasional small photobodies. Similarly, D307N, G899E exhibited either small photobodies or diffuse nuclear fluorescence.

Taken together, these observations established that most of the missense variants exhibited impairments in nuclear translocation and/or fluorescence emission (G284E, P309L, C357Y and G767R), subnuclear photobody formation (G118R, C402Y, G538D, P581S, S584F, E655K, A663V, A669V, G674D, G678E and E812K), or formation of large nuclear photobodies (R110Q, D307N, P799L, P800L, and G899E). By contrast, variants G118E, P145L and S349F exhibited both small and large nuclear photobodies, so their LOF activity may reflect impaired nuclear dynamics or their inability to interact with signaling partners like PIFs.

### YHB suppressors G767R and E812K: known loss-of-function alleles of phyB

Two prominent missense variants within the C-terminal module of phyB, G767R and E812K, have been repeatedly identified by independent screens (Wagner and Quail, 1995; Bradley et al., 1996; Chen et al., 2003). Both variants were also recovered multiple times in our screen, confirming good coverage of our mutagenesis screen and the importance of both residues for YHB signaling (Fig. 2A). These results also corroborate our recent investigation of *YHB*^*G767R*^-expressing transgenic lines, which barely complemented the long-hypocotyl phenotype of the *phyABCDE* mutant under Rc (Hu and Lagarias, 2024). The LOF activity of the G767R variant has been attributed to its lack of nuclear migration (Matsushita et al., 2003; Pfeiffer et al., 2012). YHB^G767R^ nevertheless retained limited regulatory function, presumably via its interaction with PIF3 in the cytoplasm (Hu and Lagarias, 2024). Consistent with this interpretation, the PIF3 level was reduced to some extent in the G767R variant in darkness (Fig. 2B). In addition, G767R remained signaling active in SD-grown adult plants, exhibiting strong complementation of the *phyB-5* elongated petiole and small leaf phenotypes (Fig. S6).

We next constructed *35S:YHB*^*E812K*^/*phyB-5* transgenic lines to validate the effectiveness of E812K in suppressing the constitutive activity of YHB (Fig. S7). As expected for strong suppression of YHB, all five independent *YHB*^*E812K*^ lines were nearly etiolated in the dark, almost indistinguishable from dark-grown WT and *phyB-5* seedlings (Fig. S7A). These lines exhibited reduced PIF3 levels indicating some residual activity of YHB^E812K^ in darkness (Fig. S7B). YHB^E812K^ occasionally had weak nuclear fluorescence but certainly no photobody formation (Fig. S7C). By comparison, overexpressed *YHB*^*E812K*^ was more active under Rc50, partially rescuing the long hypocotyl and small cotyledon phenotypes of *phyB-5* mutants (Fig. S7A). *YHB*^*E812K*^ overexpression also complemented the *phyB-5* adult plant phenotypes (Fig. S6). Taken together, these results indicate that both G767R and E812K variants were strong suppressors of YHB in darkness but retained sufficient phyB signaling activity in the light in both seedlings and adult plants.

### Mutations of glycine-284 suppress constitutively active phyB alleles via distinct mechanisms

We have shown that the G284E mutation restores photochemistry to YHB while rendering its dark state inactive. Previously, this residue was one of many targeted for structure-based mutagenesis (Burgie et al., 2014). In that study, the G284V variant of phyB (phyB^G284V^) exhibited a normal Pr spectrum, but was unable to fully convert to Pfr (Burgie et al., 2014). To test whether G284V influences YHB’s photochemistry, we expressed a recombinant photosensory domain construct of YHB^G284V^ in PØB chromophore-producing *E. coli* cells. Comparative analysis of recombinant YHB^G284V^ and phyB^G284V^ (as a control) revealed both proteins to be efficiently chromophorylated (Fig. 5A) and photoactive (Fig. 5B). The symmetrical photochemical difference spectra of YHB^G284V^ suggested that Pr-to-Pfr photochemistry was restored to a subpopulation of the YHB^G284V^ protein (Fig. 5B). This contrasted with the asymmetric difference spectra of phyB^G284V^ that is consistent with the formation of a bleached photoproduct. The highly similar absorption and photochemical difference spectra of YHB^G284E^ (Fig. 3B) and YHB^G284V^ (Fig. 5B) suggested that both mutations similarly influence the structure, and likely, the signaling activity of YHB.

**Fig. 5.**
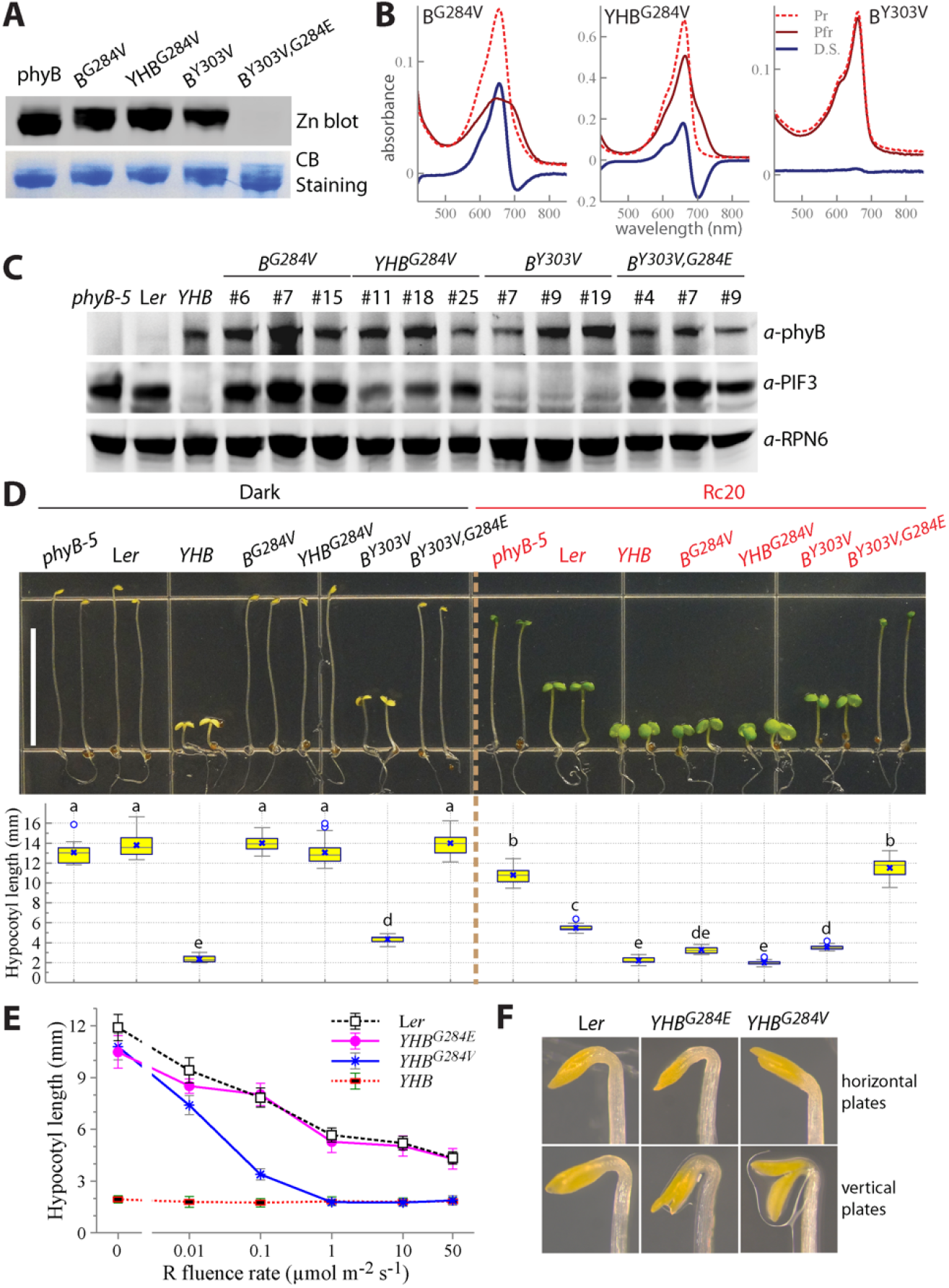
Mutations of G284 suppress the constitutive activities of YHB and phyB^Y303V^ (YVB) alleles via distinct mechanisms. (**A**) Zinc blot assay shows that phyB^Y303V,G284E^ does not covalently bind phytochromobilin. (**B**) Spectral analysis of N651 recombinant proteins phyB^G284V^, YHB^G284V^ and phyB^Y303V^. (**C**) Immunoblot assay of phyB variant and PIF3 protein levels from 4-d-old, dark-grown seedlings. (**D**) Representative 4-d-old seedlings grown in the true dark or continuous red light (Rc20) conditions (top panel), scale = 10 mm; box plots showing statistics of hypocotyl lengths, different letters indicate significant difference among groups (one-factor ANOV/Tukey’s HSD test, P < 0.001) (bottom panel), **n** = 20. (**E**) Red light fluence rate response curves, data presented as mean ± SD (**n** = 15 ∼30), the *YHB*^*G284V*^ values are from three transgenic lines #11, #18 and #25. (**F**). Apical hook phenotypes of dark-grown seedlings grown on horizontally or vertically placed plates.

To test this hypothesis, multiple transgenic lines expressing *PHYB*^*G284V*^ or *YHB*^*G284V*^ in the *phyB-5* mutant background were examined (Fig. 5C-5F). These analyses showed that the G284V mutation, like G284E (Fig. 3), strongly suppressed YHB-conferred *cop* phenotypes. However, G284V did not completely abolish YHB function *in planta*. Indeed, dark-grown *YHB*^*G284V*^ seedlings exhibited considerable PIF3 turnover, exhibited apical hook opening when seedlings were grown out of the agar medium, and displayed partially unfolded cotyledons when seedlings were grown vertically in agar medium (Fig. 5C, 5D, 5F). This residual activity differs from the fully suppressed phenotypes of dark-grown *YHB*^*G284E*^ seedlings. *YHB*^*G284V*^ seedlings were also more sensitive to red light than *YHB*^*G284E*^ seedlings as evaluated by fluence rate response curve (Fig. 5D, 5E). Indeed, *YHB*^*G284E*^ seedlings phenocopied the fluence rate response curve of L*er* wild type, while *YHB*^*G284V*^ seedlings were more strongly responsive to elevated light fluences, achieving the short hypocotyls seen for *YHB*-expressing lines at 1 µmol m^-2^ sec^-1^ and beyond (Fig. 5E).

We next tested the G284E mutation’s influence on the GOF signaling activity of another constitutively active allele of phyB. Like YHB, phyB^Y303V^ (YVB) adopts a signaling active, Pr-like state that also is photoinactive (Jeong et al., 2016; Hu and Lagarias, 2019; Choi et al., 2025). We first created multiple *35S::YVB*/*phyB-5* transgenic lines. Comprehensive phenotypic comparison, including hypocotyl elongation, cotyledon expansion and extent of growth orientation randomization, revealed that the strength of YVB’s constitutive activity is overall weaker than that of YHB (Fig. S9). We next explored whether G284E could suppress YVB function. Recombinant N651 YVB protein was well chromophorylated and spectrally locked in the Pr state (Fig. 5A, 5B) as expected from earlier studies (Jeong et al., 2016; Choi et al., 2025). By comparison, recombinant YVB^G284E^ lacked bound chromophore and red/far-red absorbance (Fig. 5A). This contrasts with the rescue of photochemistry seen for YHB^G284E^ (Fig. 3B). Due to the lack of chromophore, YVB^G284E^ also had no *in planta* function, i.e. no PIF3 protein turnover, no cop phenotypes in the dark, and no phenotypic complementation of the *phyB-5* mutant in the light (Fig. 5C, 5D). Hence, the G284E mutation acts to strongly suppress the GOF activities of YHB and YVB alleles via distinct mechanisms, i.e. rescue of WT photochemistry and loss of chromophore binding, respectively.

### Tandem P799/P800 residues are critical for YHB stability and signaling

Two tandem variants P799L and P800L within the C-terminal PAS2 domain each conferred weak ∼ moderate suppression of YHB activity (Fig. 2). Due to its sidechain forming a covalent bond with the backbone nitrogen, proline is the most rigid residue and supports formation of beta-turns in protein structures. To test whether these two tandem prolines are critical for structural stability and signaling of YHB, we expressed the YHB^P799L,P800L^ double mutant *in planta* driven by the 35S promoter. Five *YHB*^*P799L,P800L*^ transgenic lines revealed the variant protein to accumulate at levels much lower than the control germplasms *YHB, YHB*^*P799L*^ and *YHB*^*P800L*^ (Fig. 6A). This suggested that the double mutant protein was more unstable than each of the single mutants. Despite lower protein accumulation, YHB^P799L,P800L^ still effectively promoted PIF3 protein turnover in darkness. Nevertheless, *YHB*^*P799L,P800L*^ seedlings were phenotypically similar to the *phyB-5* mutant, except for a discernable cotyledon unfolding response in darkness (Fig. 6B, 6C).

**Fig. 6.**
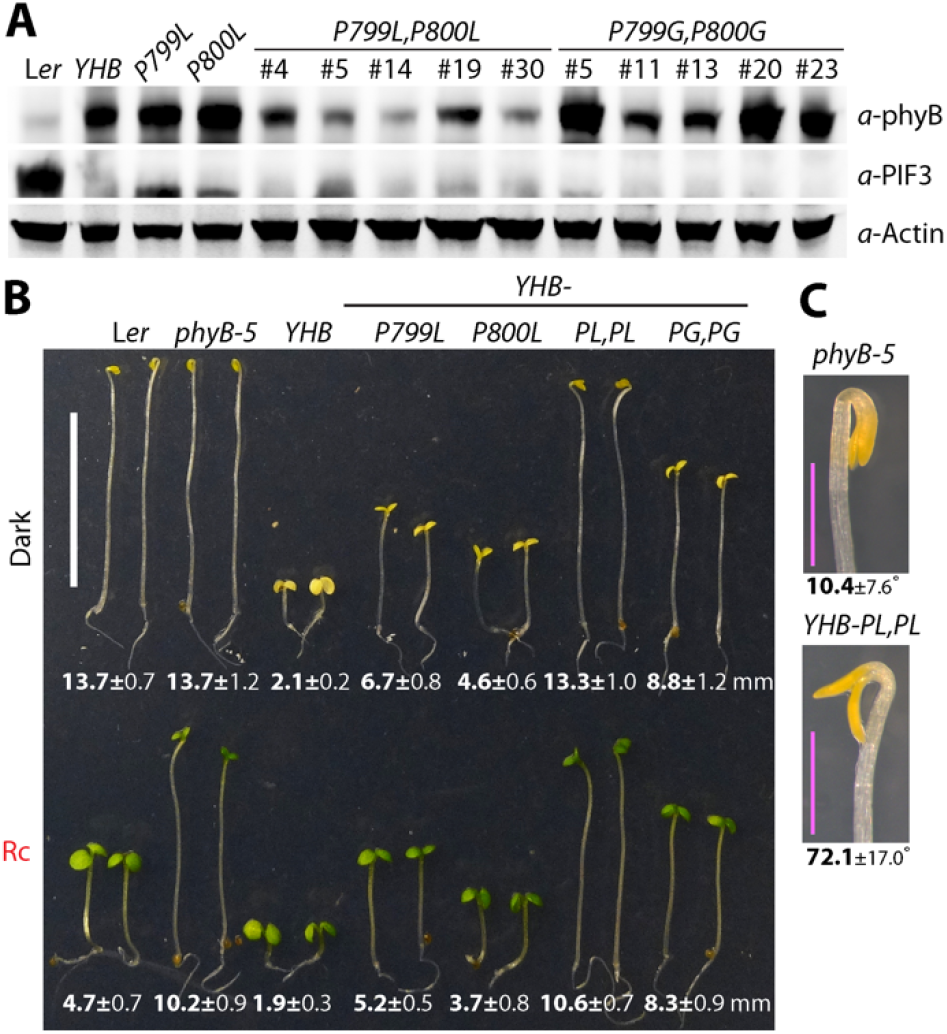
Tandem P799 and P800 residues are critical for YHB stability and signaling. (**A**) Immunoblot assays of phyB variant and PIF3 levels in dark-grown seedlings, single and double mutations were on the YHB sequence and expressed by the 35S promoter in the **phyB-5** mutant background. (**B**) Representative 4-day-old seedlings grown in the dark and under continuous red light (25 µmol m^-2^ s^-1^), hypocotyl lengths are presented as mean ± SD (**n** ≥ 15), data of *YHB*^P799L,P800L^ and *YHB*^P799G,P800G^ are from five independent lines characterized in (A), scale = 10 mm. (**C**) Dark-grown *YHB*^P799L,P800L^ seedlings marginally de-etiolate in comparison to the *phyB-5* mutant, degrees of cotyledon unfolding are presented as mean ± SD (*n* = 22 for *phyB-5* and 34 for *YHB*^*P799L,P800L*^), scale = 1 mm.

These results show that the P799L/P800L double mutation nearly abolished YHB function. In contrast to proline, glycine and alanine are the two most flexible residues. However, prolines and glycine both are statistically preferred at several beta-turn positions (Trevino et al., 2007). We therefore substituted the tandem prolines with glycines and examined the mutational consequence. In contrast to the apparent instability of YHB^P799L,P800L^, YHB^P799G,P800G^ protein accumulated to the levels comparable with YHB (Fig. 6A). Unlike the strong LOF P799L/P800L mutations, the P799G/P800G mutations moderately suppressed YHB signaling as shown by retention of dark-grown GOF cop and red light mediated hypocotyl growth suppression phenotypes in transgenic seedlings (Fig. 6B). Taken together, these results suggest that P799 and P800 play an important structural role in the formation of the signaling active state of phyB.

## Discussion

### Modularity of phyB structure and dispensability of the HKRD domain for phyB signaling

We obtained a cohort of nonsense mutations of various YHB truncation lengths. Systematic phenotypic and molecular analysis revealed that the three C-terminal nonsense alleles R1102*, Q1133* and R1136* express drastically reduced levels of truncated YHB protein, which accounts for the loss of signaling activity. Strikingly, the four nonsense alleles that lack the entire or most of the HKRD domain (Q912*, Q913*, Q921* and Q937*) exhibit more YHB signaling than those only truncating a small portion of C-terminus. These findings indicate that the HKRD is dispensable for phyB signaling, whereas the deletion of a small portion of the HKRD domain likely generates a misfolded and unstable protein that is unable to support signaling activity. These findings are consistent with the residual activity of the *phyB-28* nonsense allele in which a single guanosine nucleotide at codon 991 was deleted (Krall and Reed, 2000). The *phyB-28* protein lacks the most C-terminal 182 residues, with over two third of the HKRD domain being deleted. Accordingly, overexpression of the PHYB(1-990) fragment (similar to the *phyB-28* allele) well complemented the *phyA phyB* double mutant under red light conditions (Park et al., 2018). The concept of phyB structural modularity is additionally supported by the greater stability of nonsense mutations located between domains than those within domains. For example, the *Q784\** allele, which encodes a protein truncated between PAS1 and PAS2, yielded a much higher level of truncated YHB than *W813\** that encodes a protein truncated within the PAS2 domain. *Q784\** seedlings also had better photomorphogenic growth than *W813\** seedlings. Finally, *Q864\** encoded a protein truncated in the middle of the PAS2 domain and proved to be the only authentically null C-terminal nonsense mutation that lacked detectable YHB protein, retained WT levels of PIF3 in the dark and was phenotypically indistinguishable from the *phyB-5* mutant.

### Critical role for Glycine284

The most significant discovery from our mutagenesis screen was the G284E mutation that fully suppressed the constitutive signaling activities of Y276H (YHB) and Y303V (YVB) via distinct mechanisms. In the YHB background, the G284E mutation ablated signaling activity of YHB in the dark, while restoring its ability to generate a signaling active Pfr state in the light. Hence, this mutation generated a photoreceptor that functions similarly to WT phyB - representing a true phenotypic reversion of the *YHB* allele. By comparison, G284E blocked chromophorylation of YVB, nullifying the function of YVB^G284E^ regardless of light conditions. Interestingly, the G284E mutation was previously identified in a genetic screen as a chromophore-binding LOF allele which conferred long hypocotyls under Rc (Oka et al., 2008). Taken together, these studies show that the G284E mutation suppresses the signaling activities of phyB, YHB and YVB proteins by distinct mechanisms.

The context-dependent phenotypes of the G284E variant lines suggest a structural interaction among G284, Y/H276 and Y303. Indeed, all of these residues are localized within 3-5.7 Å from each other within the chromophore binding pocket (Fig. 7, (Li et al., 2022; Wang et al., 2024; Jia et al., 2025)). In WT phyB, the *Z*-to-*E* photoisomerization of chromophore leads to a repositioning of the phenolic sidechains of Y276 and Y303 into the gap above G284 (Fig. 7A, 7B). In Pr, the phenol sidechain of Y276 adopts an inward facing orientation above the gap while Y303 is pointing outward. In Pfr and YHB (Wang et al., 2024), Y303 reorients inward excluding Y276 (or YHB) which adopts a rotamer perpendicular to the plane of the bilin chromophore. Replacement of G284 with glutamate inserts a large residue with a negative charge within the chromophore binding pocket that would prevent repositioning of Y276 and Y303 phenolic sidechains into the gap. Indeed, replacement of G284 with glutamate and reoptimizing the rotamers of E284, H276 and Y303 sidechains *in silico* generates an energy-minimized structure in which the phenolic sidechain of Y303 adopts an outward oriented geometry similar with that seen in Pr (Fig. 7C). H276 instead adopts an orientation that is perpendicular to the plane of the chromophore similar to the geometry seen in Pfr. Because G284V had a similar though slightly weaker effect than G284E in suppressing YHB, we speculate that substituting the glycine with any other residue with a bulky sidechain would render similar mutational impact.

**Fig. 7.**
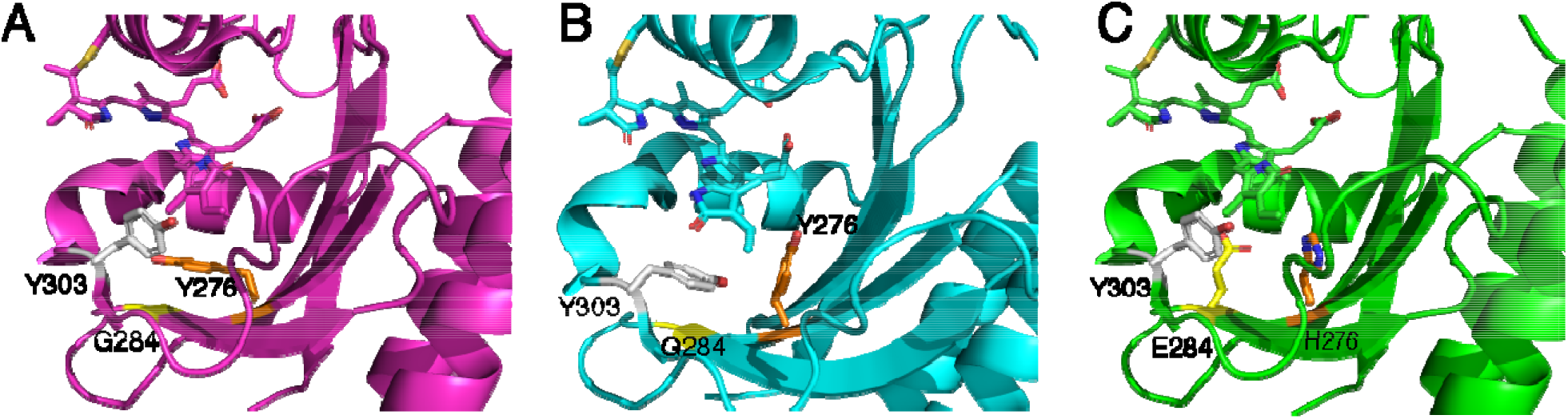
Replacement of G284 with glutamate in YHB prevents reorientation of Y303 that accompanies Pfr formation. Structures of the chromophore binding pockets of wild-type Arabidopsis phyB **(A)** as Pr (PDB 4OUR, Burgie et al., 2014) and **(B)** as Pfr (PDB 8YB4, Wang et al., 2024) which document the tyrosine exchang mechanism that occurs upon Pr-to-Pfr photoconversion. In Pr, the phenol sidechain of Y276 (orange) adopts an inward facing orientation into the gap above G284 (yellow) while Y303 (white) is pointing outward. In Pfr and also in YHB (Wang et al., 2024), Y303 reorients inward to occupy the gap above G284 while excluding Y276 whic orients perpendicular to the plane of the bilin chromophore. (C) Model of the YHB^G284E^ double mutant chromophore binding site created by replacement of G284 with glutamate in the Pr structure and selectin sidechain rotamers for Y303 and H276 which minimize steric clashes. This prevents inward orientation of either tyrosine sidechain, both of which are oriented away from the chromophore. Structures are rendered with PyMOL (https://www.pymol.org/).

Whereas the modeled structure of YHB^G284E^ exhibits features of both Pr and Pfr (Fig. 7C), signal transmission is mediated via interactions of the loop region in which Y303 is located with the tongue insert region of the PHY domain. During Pr-to-Pfr conversion, this tongue insert refolds from a beta hairpin to an alpha helix - a process which involves reorientation of Y303 (Li et al., 2022; Wang et al., 2024; Jia et al., 2025). As the Y303 orientation in YHB^G284E^ is similar to that found in the inactive Pr state (compare Figs. 7A and 7C), this could account for the suppression of YHB’s constitutive activity by the G284E mutation. The inability to form the signaling active Pfr state by Y303 repositioning is likely also responsible for the LOF activity of the phyB^G284E^ allele (Oka et al., 2008). The importance of Y303 orientation probably also accounts for the GOF activity of YVB in which Y303 is substituted with valine (Jeong et al., 2016; Choi et al., 2025). It is reasonable to assume that V303 in this variant favors an inward facing Pfr-like active conformation of the V303 sidechain. The surprising ability of YHB^G284E^ to regain photoactivity remains a mystery at present. While the proposed structure of YHB^G284E^ would allow for the chromophore to undergo Z-to-E photoisomerization (Fig. 7C), it is hard to rationalize how the inward rotation of Y303 might occur given the steric hindrance of the E284 sidechain. We suspect that the chromophore pocket is remodeled to accommodate this reorientation. This open question will require additional structural work to fully resolve.

### P309L ablates YHB activity

In contrast to the G284E mutation that restores R/FR physiological function to YHB, P309L ablates YHB function regardless of light conditions. The abnormal spectra plus reduced chromophorylation level of the YHB^P309L^ variant indicate that Y276H-elicited conformational activation of phyB signaling was blocked by the P309L mutation. Consistent with this result, the P309L mutation was previously reported to reduce chromophorylation and inhibit phyB activity, although it did not render an abnormal spectra as seen for YHB^P309L^ (Oka et al., 2008). Hence, P309 also plays an important role in phyB photoactivation. Interestingly, the neighboring D307N mutation also imparts a secondary absorption band in the yellow-orange region, similar to the effects of P309L and C357Y. Compared with the full phenotypic suppression by the P309L mutation, D307N only weakly suppressed the GOF activity of YHB. However, the D307A mutation markedly prevented Pr-to-Pfr photoconversion of recombinant phyB protein and strongly compromised phyB signaling *in planta* (Zhang et al., 2013; Burgie et al., 2014). The D307 residue lies within the invariant phyB-D_307_I_308_P_309_ triplet motif and has been proposed to be critical for the protonation/deprotonation cycle of the chromophore that occurs during photoactivation (Burgie et al., 2014). Substitutions effecting chromophore pK values (and therefore on protonation of the static photostate) are known in Cph1 and can induce absorption in the same spectral window, so P309 could function as a rigid residue to provide a structural framework to support D307 in coordinating chromophore placement and photoisomerization.

### Protein stability

Two missense mutations G118R and C402Y greatly reduce YHB abundance, likely through decreased stability. Previous studies have shown that the PHYB^G118R^-GFP variant accumulates to a lower protein abundance than the PHYB-GFP control (Chen et al., 2003). By comparison, C402Y elicits YHB instability more profoundly than G118R. Recombinant G118R and C402Y variants as well as the G538D variant also poorly bind chromophore and exhibit distinct patterns of protein size and zinc blot by comparison with C357Y - a mutation that eliminates covalent chromophore binding. We propose that the G118R, C402Y and G538D mutations alter the conformational state of the chromophore docking pocket to disfavor chromophore binding to C357. Within the regulatory module region, the P799L,P800L double mutation was also found to lead to protein instability. Structural rigidity inherently provided by these tandem prolines is likely essential for proper folding and stabilizing YHB/phyB protein. The role of such structural rigidity in coordinating (photo)activation of YHB/phyB signaling awaits further investigation.

### Holistic reflections on missense variants

Our large-scale EMS mutagenesis screen attained good coverage of key residues for YHB/phyB signaling, exemplified by the five strong variants G284E, P309L, S584F, G767R and E812K which have also been repeatedly identified by earlier genetic screens (Wagner and Quail, 1995; Bradley et al., 1996; Chen et al., 2003; Oka et al., 2008). We also identified 14 novel variants, including two strong variants C402Y and G538D, both of which are poorly chromophorylated. A significant number of variants within the photosensory module exhibited reduced chromophorylation, corroborating the indispensability of bound bilin for the YHB function (Su and Lagarias, 2007). Surprisingly, mutations in C357, the conserved residue for covalent binding of the PØB chromophore, have not been previously identified by mutagenesis screens. Besides the present study, the cysteine-to-tyrosine mutation has only been reported for the wheat phyC^B^ equivalent C323Y, which had no chromophore absorption as expected for a phy protein lacking bound bilin (Chen et al., 2014). By contrast, the Arabidopsis YHB^C357Y^ variant strongly binds bilin noncovalently and displays significant visible light absorption. Despite its non-covalent binding of chromophore and weak photoactivity, YHB^C357Y^ is a strong LOF allele. Consistently, YHB^C357A^-GFP was reported to also bind chromophore non-covalently, but it unexpectedly conferred discernable cop phenotypes in transgenic plants (Oka et al., 2011). The substitution type for C357 thus seems to have varied effect in suppressing the YHB signaling activity.

In Table S2, we assemble all known phyB missense LOF and GOF variants identified by EMS mutagenesis screens and/or created by structure-informed mutagenesis - most of which have been functionally assessed *in planta*. Many of these variants, especially those from structure-informed mutagenesis, were missed by our screen, partially because EMS effects exclusive G/C-to-A/T conversions which limits residue alteration. On the other hand, YHB is photochemically inert, thus mutations merely affecting photoconversion and/or thermal (dark) reversion would not be expected to be identified. One notable exception was S584F which profoundly accelerates phyB dark reversion (Oka et al., 2008; Burgie et al., 2014; Jeon et al., 2024; Wang et al., 2024). S584 is localized in the PHY tongue region that interacts with the chromophore associated loop region in which Y303 is found. As such, it is in a good position to stabilize the activated alpha helical tongue region in the Pfr form. Replacement of S584 with phenylalanine likely destabilizes the alpha helical ‘active’ conformer that is seen in the YHB (Wang et al., 2024; Jia et al., 2025), leading to its loss-of-function. By contrast, mutations affecting YHB nuclear import, photobody formation and interaction with downstream signaling partners are particularly enriched by current screen endeavor.

## Materials and Methods

### Seedling growth conditions and phenotypic analysis

Arabidopsis seeds were surface sterilized with 75% ethanol for ∼12 min, resuspended with 0.1% phytagar solution, and sown on half-strength Murashige and Skoog□(MS) media (pH 5.7) solidified with 0.8% phytagar. After three or four days of stratification, seeds were exposed to□3 h white light (∼□80 µmol m^−2^ s^−1^) to induce synchronized germination. Seedlings were then grown in the dark, in the true dark (Leivar et al., 2008), or under continuous red light of indicated fluence rate. Seedlings were photographed and measured digitally using the NIH ImageJ software (https://imagej.net/ij/).

### Single-insertion *35S::eYHB*/*phyB-5* lines and EMS mutagenesis screen of YHB suppressor mutants

The synthetic eYHB-3xFLAG fragment, in which nearly all restriction sites of the YHB sequence were synonymously mutated (see Fig. S3) (Hu and Lagarias, 2017), were subcloned into the pJM61 binary vector through *Kpn*I/*Xba*I ligation (Su and Lagarias, 2007). The resultant pJM61-35S::eYHB-3xFLAG construct was transformed into the *phyB-5* mutant by the agrobacterium-mediated floral dip method (Clough and Bent, 1998). Two authentic single-insertion transgenic lines #3 and #4 identified from 15 lines (Fig. S1) were used for ethyl methane sulfonate (EMS) mutagenesis. Approximately 50,000 seeds in total from two lines were treated with 0.2∼0.3% (v/v) EMS for 16 hours with gentle shaking. After thorough rinse with water multiple times, these M1 seeds were resuspended with 0.1% phytagar solution and sown on the soil for growth. M2 seeds harvested from multiple M1 plants of the same growth pot were pooled as one mutation family and proceeded to dark growth screen to identify etiolated seedlings or dampened *cop* seedlings showing significantly elongated hypocotyls. Intermittent low light was given on day 2 and day 3 to prevent photodamage on moderate or weak cop seedlings once transitioned to ambient light on day 4 (Fig. S2). M3 lines were grown under different light conditions and were also backcrossed with the *phyB-5* mutant to identify intragenic mutants, from which the *eYHB-3xFLAG* transgene was PCR amplified (LongAmp Taq DNA polymerase, New England Biolabs) (Hu and Lagarias, 2020) and sequenced to determine nucleotide mutations (Fig. S2). Mutations identified from multiple independent M2 plants of the same seed family pool were not regarded as independent. Some lines such as R110Q, P309L and C402Y bearing traits irrelevant to phyB signaling were further backcrossed with *phyB-5* to segregate out the unrelated mutations.

### Construction of transgenic plants

The *PHYB*^*G284V*^, *YHB*^*G284V*^, *PHYB*^*Y303V*^, *PHYB*^*Y303V,G284E*^, *YHB*^*E812K*^, *YHB*^*P799L,P800L*^ and *YHB*^*P799G,P800G*^ mutation variants were created by site-directed mutagenesis on the pUC57-*ePHYB-3xFLAG* or pUC57-*eYHB-3xFLAG* constructs (Hu and Lagarias, 2017), and then subcloned into the pJM61 binary vector through *Kpn*I/*Xba*I ligation. These *PHYB*/*YHB* mutant overexpression constructs driven by the 35S promoter were transformed into the *phyB-5* mutant by the agrobacterium-mediated floral dip method (Clough and Bent, 1998). Multiple homozygous, genetically single-insertion lines for each construct were obtained through standard genetic procedures for phenotypic analyses.

### Recombinant protein preparation and spectra analysis of N651 variants

The partial fragments of *eYHB* variants from individual suppressor lines were amplified using the primers oWH202 and PHYB-R3 (Table S3), which then served as the template for the secondary PCR using the primers oWH295 and oWH314 (Table S3) to amplify the N-terminal photosensory module sequences (N651). Wild-type *ePHYB* and variant sequences created by site-directed mutagenesis directly served as the template to amplify the N651 fragments using oWH295 and oWH314. These N651 fragments were subcloned into the pBAD-CBD vector through *Nco*I/*Xma*I ligation. The resultant pBAD-YHB/PHYB-CBD constructs were transformed into the *E. coli* strain LMG194 (Invitrogen) hosting the pPL-PØB phytochromobilin biosynthetic expression plasmid (Gambetta and Lagarias, 2001; Fischer et al., 2005). Such obtained dual ampicillin- and kanamycin-resistant transformants were cultured and induced for protein expression as previously described (Fischer et al., 2005). Collected cell pellets were resuspended in lysis buffer (20 mM Na-HEPES pH 8.0, 500 mM NaCl, 1 mM EDTA, 0.1% Triton X-100) and homogenized using a microfluidizer (Microfluidics, model #Y110Y).

Homogenates were centrifuged 35,00 rpm x 30 min. The resultant lysate supernatants were loaded at a rate of 1∼1.5 ml/min onto chitin resin column (NEB, #S6651) that was pre-equilibrated with 10x bed volumes of wash buffer (20 mM Na-HEPES pH 8.0, 500 mM NaCl, 1 mM EDTA) under green light in cold room. The column was then washed with 10x bed volumes of wash buffer. Bound recombinant protein was released from the column by incubation overnight with cleavage buffer (wash buffer + 50 mM DDT, freshly prepared) to cleave off the intein-CBD tag and eluted with cleavage buffer. The collected eluate was then dialyzed against the dialysis buffer (25 mM TES-KOH pH 8.0, 25 mM KCl, 10% glycerol) (first liter, three hours; second liter, overnight). Spectrophotometric measurements of purified recombinant proteins were performed using a Cary 50 UV-visible spectrophotometer as described previously (Fischer et al., 2005). Amicon® Ultra-0.5 10k centrifugal filter devices were used to concentrate protein solutions as needed.

### SDS PAGE and Zinc Blot assays

Zinc blot assay was as described previously (Berkelman and Lagarias, 1986) with appropriate modifications below. Purified recombinant proteins were quantified by BCA protein assay (Pierce), then mixed with 4x SDS sample loading buffer at room temperature and loaded for SDS-PAGE. After electrophoresis, proteins were electroblotted onto Immobilon®-FL PVDF membrane (Millipore). The membrane was incubated with 1.3M zinc acetate solution for 30 min and rinsed with distilled water three times afterwards. LI-COR 9120 Odyssey Infrared Imaging System was employed to scan the membrane using the 685 nm laser channel to capture the fluorescence emitted by biliprotein-zinc acetate complex.

### Protein extraction and immunoblot assay

Dark-grown, 4-day-old seedlings were harvested under dim green light conditions, ground to fine powder in liquid nitrogen, and used to extract protein following an established protocol (Su and Lagarias, 2007). Immunoblot assay procedures were as described previously (Jones et al., 2015; Hu and Lagarias, 2017). The B1 mAb recognizing an epitope between residues 90 and 103 of phyB (Somers et al., 1991; Wagner et al., 1996) was used to immunodetect phyB, YHB and truncated YHB variants (1:500 dilution). Polyclonal anti-PIF3 (Chen et al., 2010), monoclonal anti-Actin (#MA1-744, Thermo Scientific), monoclonal anti-Alpha Tubulin (#MA1-19162, Thermo Scientific), and polyclonal anti-RPN6 (#BML-PW8370, Enzo Life Sciences) antibodies were used at 1:1000 dilution.

### Quantitative RT-PCR and confocal fluorescence microscopy

Total RNA was extracted from dark-grown, four-day-old seedlings using RNeasy Plant Mini Kit (Qiagen). Reverse transcription and real-time quantitative PCR procedures were performed as described previously (Hu et al., 2020). Primers used for PCR are list in Table S3. Confocal fluorescence microscopy was performed as described previously (Su and Lagarias, 2007).

## Accession Numbers

Sequence data from this article can be found in the Arabidopsis Information Resource (www.arabidopsis.org) under the following accession numbers: At2g18790 (*PHYB*), At1g09530 (*PIF3*), At2g31380 (*STH*), At5g54190 (*PORA*), and At1g29150 (RPN6).

## Acknowledgements

We thank Shelley Martin for training of expression and purification of recombinant proteins. We also thank Prof. Peter Quail and Meng Chen for the phyB and PIF3 antibodies, respectively. This work was supported in part by the National Institutes of Health grants RO1:GM068552 and R35GM139598 awarded to JCL.

## List of Supplementary Data files

**Figure S1**. Preparation of authentic single-insertion *35S::eYHB-3xFLAG/phyB-5* lines for EMS mutagenesis screen.

**Figure S2**. Possible mutations suppressing the YHB function and the workflow of isolating and identifying intragenic YHB suppressor mutants.

**Figure S3**. The eYHB DNA and protein sequences and residues (colored and bold) for potential nonsense mutation by EMS mutagenesis.

**Figure S4**. Adult plants of C-terminal nonsense mutations grown under short-day conditions for 36 days.

**Figure S5**. Reduction of steady YHB protein levels by the C402Y mutation is likely due to protein instability, not by reduced transcription levels.

**Figure S6**. Missense suppressor plants grown under short-day conditions for 40 days.

**Figure S7**. Validation of E812K suppressive effect on the YHB function by independent transgenic lines.

**Figure S8**. Confocal microscopy examination of subnuclear distributions of missense variants from dark-grown seedlings.

**Figure S9**. The constitutive activity of phyB^Y303V^ (YVB) is weaker than phyB^Y276H^ (YHB).

**Table S1**. Identified mutation alleles in this study.

**Table S2**. PHYB missense mutations identified or characterized from previous and current work (in excel file).

**Table S3**. Primers used in the current study.

