## supplementary materials for "Intragenic suppressor screen of YHB identifies novel and known loss-of-function alleles of Arabidopsis phytochrome B"

**Supplementary Information**

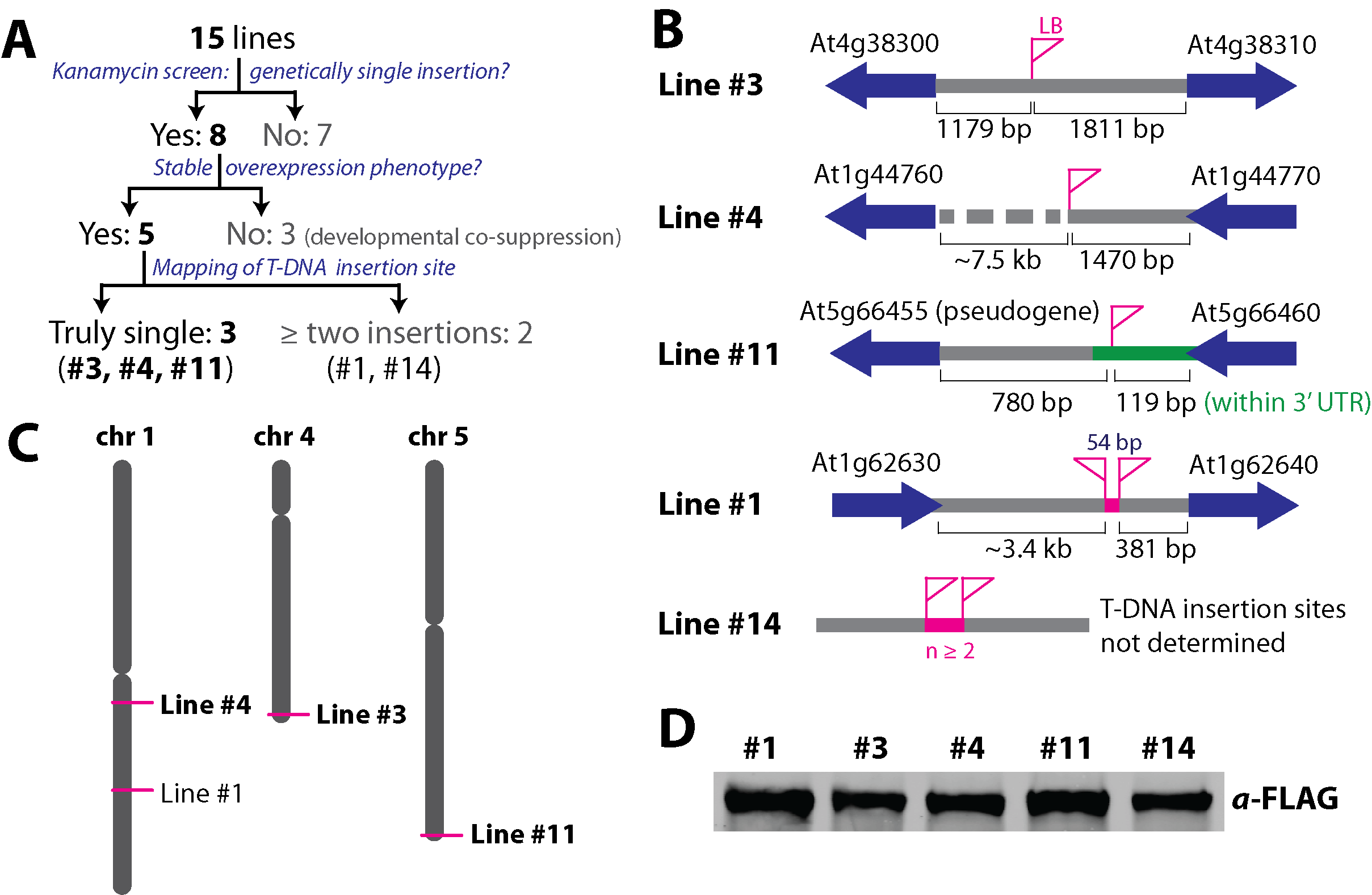

**Fig. S1 Preparation of authentic single-insertion *35S::eYHB-3xFLAG/phyB-5* lines for EMS mutagenesis screen.** (**A**) The workflow of obtaining three authentic single-insertion transgenic lines. (**B**) Schematic diagrams of T-DNA insertion locations and insertion copies of five stable, genetically single-insertion lines. (**C**) Chromosomal positions of T-DNA insertion sites of four stable, genetically single-insertion lines. (**D**) Immunoblot assay confirms similar eYHB-3xFLAG expression levels of the five genetically single-insertion lines.

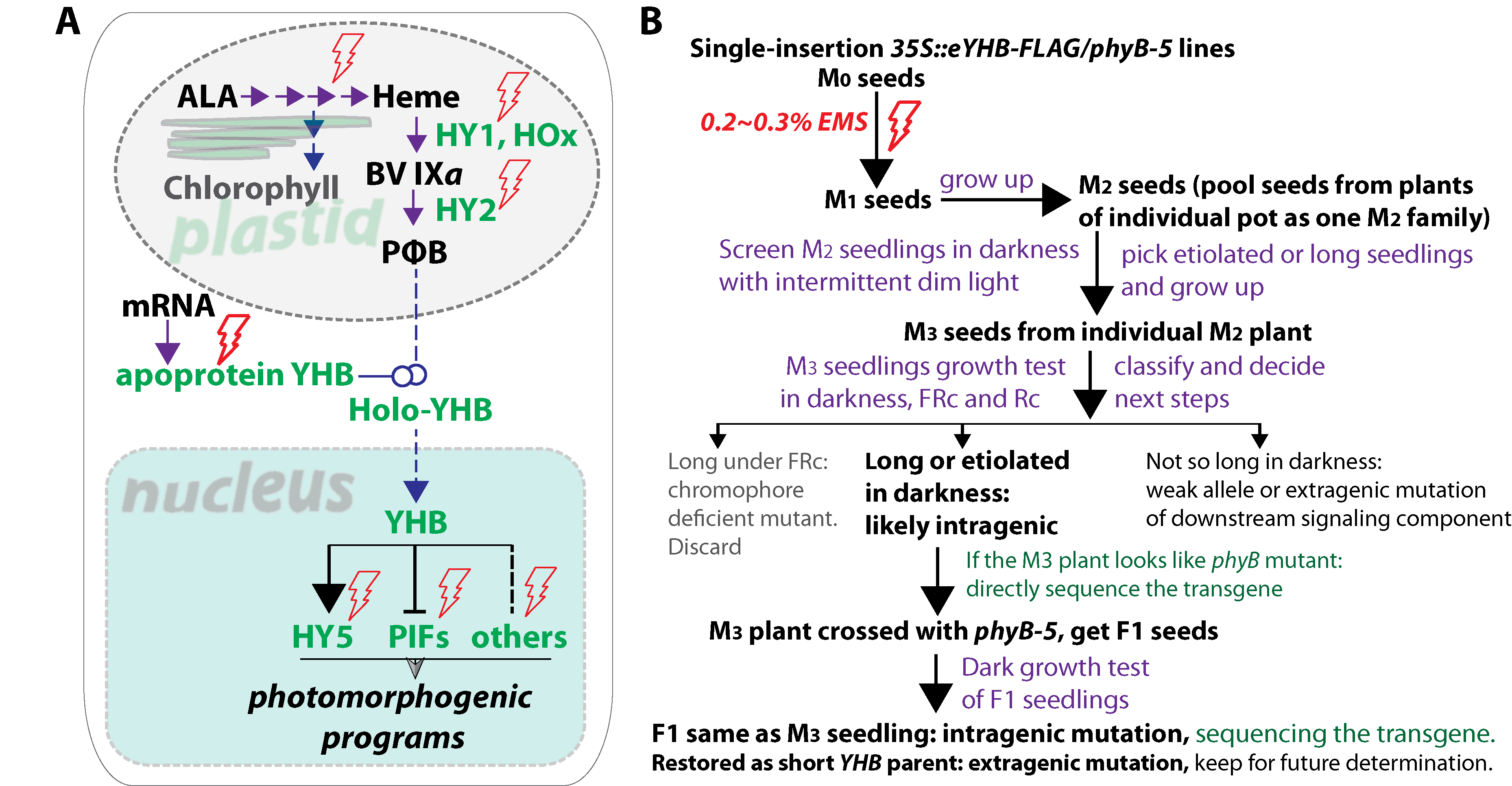

**Fig. S2 Possible mutations suppressing the YHB function and the workflow of isolating and identifying intragenic YHB suppressor mutants.** (**A**) Simplified YHB signaling pathway illustrating potential targets of suppressor mutations (denoted by lightning symbols): chromophore biosynthesis genes, the YHB transgene itself, and downstream signaling components. (**B**) The workflow of EMS mutagenesis and screening for intragenic suppressor mutants.

```
   1 ATGGTTTCCGGAGTCGGGGGTAGTGGCGGTGGCCGTGGCGGTGGCCGTGGCGGAGAAGAAGAACCGTCGTCAAGTCACACTCCTAATAACCGAAGAGGAGGAGAACAAGCTCAATCGTCGGGAACGAAATCTCTCAGACCAAGAAGCAAC
   1  M  V  S  G  V  G  G  S  G  G  G  R  G  G  G  R  G  G  E  E  E  P  S  S  S  H  T  P  N  N  R  R  G  G  E  Q  A  Q  S  S  G  T  K  S  L  R  P  R  S  N

 151 ACTGAATCAATGAGCAAAGCAATTCAACAGTACACCGTtGACGCAAGACTCCACGCCGTTTTCGAACAATCCGGCGAATCAGGGAAATCATTCGACTACTCACAATCACTCAAAACGACGACGTACGGTTCCTCTGTACCTGAGCAACAG
  51  T  E  S  M  S  K  A  I  Q  Q  Y  T  V  D  A  R  L  H  A  V  F  E  Q  S  G  E  S  G  K  S  F  D  Y  S  Q  S  L  K  T  T  T  Y  G  S  S  V  P  E  Q  Q

 301 ATCACAGCTTATCTCTCTCGAATCCAGCGAGGTGGTTACATTCAGCCTTTCGGATGTATGATCGCCGTCGATGAATCCAGTTTCCGGATCATCGGTTACAGTGAAAACGCCAGAGAAATGTTAGGGATTATGCCTCAATCTGTTCCTACT
 101  I  T  A  Y  L  S  R  I  Q  R  G  G  Y  I  Q  P  F  G  C  M  I  A  V  D  E  S  S  F  R  I  I  G  Y  S  E  N  A  R  E  M  L  G  I  M  P  Q  S  V  P  T

 451 CTTGAGAAACCTGAGATTCTAGCTATGGGAACTGATGTGcGATCTTTGTTCACTTCTTCGAGtTCGATTCTACTCGAaCGTGCTTTCGTTGCTaGAGAGATTACCTTGTTAAATCCGGTTTGGATtCATTCCAAGAATACTGGTAAACCG
 151  L  E  K  P  E  I  L  A  M  G  T  D  V  R  S  L  F  T  S  S  S  S  I  L  L  E  R  A  F  V  A  R  E  I  T  L  L  N  P  V  W  I  H  S  K  N  T  G  K  P

 601 TTTTACGCCATTCTTCATAGGATTGATGTTGGTGTTGTTATTGATTTAGAGCCAGCTAGAACTGAAGATCCTGCGCTTTCTATTGCTGGTGCTGTTCAATCGCAGAAACTCGCGGTTCGTGCGATTTCTCAGTTACAGGCTCTTCCTGGT
 201  F  Y  A  I  L  H  R  I  D  V  G  V  V  I  D  L  E  P  A  R  T  E  D  P  A  L  S  I  A  G  A  V  Q  S  Q  K  L  A  V  R  A  I  S  Q  L  Q  A  L  P  G

 751 GGAGATATTAAGCTgTTGTGTGACACTGTCGTGGAAAGTGTGAGGGACTTGACTGGTTATGATCGTGTTATGGTTCATAAGTTTCATGAAGATGAGCATGGAGAAGTTGTAGCTGAGAGTAAACGAGACGATTTAGAGCCTTATATTGGA
 251  G  D  I  K  L  L  C  D  T  V  V  E  S  V  R  D  L  T  G  Y  D  R  V  M  V  H  K  F  H  E  D  E  H  G  E  V  V  A  E  S  K  R  D  D  L  E  P  Y  I  G

 901 CTGCATTATCCTGCTACTGATATTCCTCAAGCGTCAAGGTTCTTGTTTAAGCAGAACCGTGTCCGAATGATAGTAGATTGCAATGCCACACCTGTTCTTGTGGTCCAGGACGATAGGCTAACTCAGTCTATGTGCTTGGTTGGTTCTACT
 301  L  H  Y  P  A  T  D  I  P  Q  A  S  R  F  L  F  K  Q  N  R  V  R  M  I  V  D  C  N  A  T  P  V  L  V  V  Q  D  D  R  L  T  Q  S  M  C  L  V  G  S  T

1051 CTTAGGGCTCCTCATGGTTGTCACTCTCAGTATATGGCTAACATGGGATCTATTGCGTCTTTAGCAATGGCGGTTATtATCAATGGAAATGAAGATGATGGGAGCAATGTAGCTAGTGGAAGAAGCTCGATGAGGCTTTGGGGTTTGGTT
 351  L  R  A  P  H  G  C  H  S  Q  Y  M  A  N  M  G  S  I  A  S  L  A  M  A  V  I  I  N  G  N  E  D  D  G  S  N  V  A  S  G  R  S  S  M  R  L  W  G  L  V

1201 GTTTGCCATCACACTTCTTCTCGCTGCATACCGTTTCCGCTAAGGTATGCTTGTGAGTTTTTGATGCAGGCTTTCGGTTTACAGTTAAACATGGAATTGCAGTTAGCTTTGCAAATGTCAGAGAAACGCGTTTTGAGAACGCAGACACTG
 401  V  C  H  H  T  S  S  R  C  I  P  F  P  L  R  Y  A  C  E  F  L  M  Q  A  F  G  L  Q  L  N  M  E  L  Q  L  A  L  Q  M  S  E  K  R  V  L  R  T  Q  T  L

1351 TTATGTGATATGCTTCTGCGTGACTCGCCTGCTGGAATTGTTACACAGAGTCCCAGTATCATGGACTTAGTGAAATGTGACGGTGCAGCATTTCTTTACCACGGGAAGTATTACCCGTTGGGTGTTGCTCCTAGTGAAGTTCAGATAAAA
 451  L  C  D  M  L  L  R  D  S  P  A  G  I  V  T  Q  S  P  S  I  M  D  L  V  K  C  D  G  A  A  F  L  Y  H  G  K  Y  Y  P  L  G  V  A  P  S  E  V  Q  I  K

1501 GATGTTGTGGAGTGGTTGCTTGCGAATCATGCGGATTCAACCGGATTAAGCACTGATAGTTTAGGCGATGCGGGGTATCCCGGTGCAGCgGCGTTAGGGGATGCTGTGTGCGGTATGGCAGTTGCATATATCACAAAAAGAGACTTTCTT
 501  D  V  V  E  W  L  L  A  N  H  A  D  S  T  G  L  S  T  D  S  L  G  D  A  G  Y  P  G  A  A  A  L  G  D  A  V  C  G  M  A  V  A  Y  I  T  K  R  D  F  L

1651 TTTTGGTTTCGATCTCACACTGCGAAAGAAATCAAATGGGGAGGCGCTAAGCATCATCCGGAGGATAAAGATGATGGGCAACGAATGCAcCCTCGTTCGTCCTTTCAGGCTTTTCTTGAAGTTGTTAAGAGCCGGAGTCAGCCtTGGGAA
 551  F  W  F  R  S  H  T  A  K  E  I  K  W  G  G  A  K  H  H  P  E  D  K  D  D  G  Q  R  M  H  P  R  S  S  F  Q  A  F  L  E  V  V  K  S  R  S  Q  P  W  E

1801 ACTGCGGAAATGGATGCGATTCACTCGCTCCAGCTTATTCTGAGAGACTCTTTcAAAGAATCTGAGGCGGCTATGAACTCTAAAGTTGTGGATGGTGTGGTTCAGCCATGTAGGGATATGGCGGGGGAACAGGGGATTGATGAGTTAGGT
 601  T  A  E  M  D  A  I  H  S  L  Q  L  I  L  R  D  S  F  K  E  S  E  A  A  M  N  S  K  V  V  D  G  V  V  Q  P  C  R  D  M  A  G  E  Q  G  I  D  E  L  G

1951 GCAGTTGCAAGAGAGATGGTTAGGCTCATTGAGACTGCAACTGTTCCTATATTCGCTGTGGATGCCGGAGGCTGCATCAATGGATGGAACGCTAAGATTGCAGAGTTGACAGGTCTCTCAGTTGAAGAAGCTATGGGGAAGTCTCTGGTT
 651  A  V  A  R  E  M  V  R  L  I  E  T  A  T  V  P  I  F  A  V  D  A  G  G  C  I  N  G  W  N  A  K  I  A  E  L  T  G  L  S  V  E  E  A  M  G  K  S  L  V

2101 TCTGATTTAATATACAAAGAGAATGAAGCAACTGTCAATAAGCTgCTTTCTCGTGCTTTGAGAGGGGACGAGGAAAAGAATGTGGAGGTTAAGCTGAAAACTTTCAGCCCCGAACTACAAGGGAAAGCAGTTTTTGTGGTTGTGAATGCT
 701  S  D  L  I  Y  K  E  N  E  A  T  V  N  K  L  L  S  R  A  L  R  G  D  E  E  K  N  V  E  V  K  L  K  T  F  S  P  E  L  Q  G  K  A  V  F  V  V  V  N  A

2251 TGTTCCAGCAAGGACTACTTGAACAACATTGTCGGCGTTTGTTTTGTTGGACAAGACGTTACaAGTCAGAAAATCGTAATGGATAAGTTCATCAACATACAAGGAGATTACAAGGCTATTGTACATAGCCCAAACCCTCTAATCCCGCCA
 751  C  S  S  K  D  Y  L  N  N  I  V  G  V  C  F  V  G  Q  D  V  T  S  Q  K  I  V  M  D  K  F  I  N  I  Q  G  D  Y  K  A  I  V  H  S  P  N  P  L  I  P  P

2401 ATTTTTGCTGCTGACGAGAACACtTGCTGCCTGGAATGGAACATGGCGATGGAAAAGCTgACGGGTTGGTCTCGCAGTGAAGTGATTGGGAAAATGATTGTCGGGGAAGTGTTTGGGAGCTGTTGCATGtTAAAGGGTCCTGATGCTTTA
 801  I  F  A  A  D  E  N  T  C  C  L  E  W  N  M  A  M  E  K  L  T  G  W  S  R  S  E  V  I  G  K  M  I  V  G  E  V  F  G  S  C  C  M  L  K  G  P  D  A  L

2551 ACCAAGTTCATGATTGTATTGCATAATGCGATTGGTGGtCAAGATACGGATAAGTTCCCTTTCCCATTCTTTGACCGCAATGGGAAGTTTGTTCAGGCTCTATTGACTGCAAACAAGCGGGTTAGCCTCGAaGGAAAGGTTATTGGGGCT
 851  T  K  F  M  I  V  L  H  N  A  I  G  G  Q  D  T  D  K  F  P  F  P  F  F  D  R  N  G  K  F  V  Q  A  L  L  T  A  N  K  R  V  S  L  E  G  K  V  I  G  A

2701 TTCTGTTTCTTGCAAATCCCGAGCCCTGAGtTGCAGCAAGCaTTAGCAGTCCAACGGAGGCAGGACACAGAGTGTTTCACGAAGGCAAAAGAGTTGGCTTATATTTGTCAGGTGATAAAGAATCCTTTGAGCGGTATGCGTTTCGCAAAC
 901  F  C  F  L  Q  I  P  S  P  E  L  Q  Q  A  L  A  V  Q  R  R  Q  D  T  E  C  F  T  K  A  K  E  L  A  Y  I  C  Q  V  I  K  N  P  L  S  G  M  R  F  A  N

2851 TCATTGTTGGAGGCCACAGACTTGAACGAGGACCAGAAGCAGTTACTTGAAACAAGTGTTTCTTGCGAGAAACAGATtTCAAGGATCGTCGGGGACATGGATCTTGAAAGCATTGAAGACGGTTCATTTGTGCTAAAGAGGGAAGAGTTT
 951  S  L  L  E  A  T  D  L  N  E  D  Q  K  Q  L  L  E  T  S  V  S  C  E  K  Q  I  S  R  I  V  G  D  M  D  L  E  S  I  E  D  G  S  F  V  L  K  R  E  E  F

3001 TTCCTTGGAAGTGTCATAAACGCGATTGTAAGTCAAGCGATGTTCTTATTAAGGGACAGAGGTCTTCAaCTGATCCGTGACATTCCCGAAGAGATCAAATCAATAGAGGTTTTTGGAGACCAGATAAGGATTCAACAGCTCCTGGCTGAG
1001  F  L  G  S  V  I  N  A  I  V  S  Q  A  M  F  L  L  R  D  R  G  L  Q  L  I  R  D  I  P  E  E  I  K  S  I  E  V  F  G  D  Q  I  R  I  Q  Q  L  L  A  E

3151 TTTCTGCTGAGTATAATCCGGTATGCACCATCTCAAGAGTGGGTGGAGATCCATTTAAGCCAACTTTCAAAGCAAATGGCTGATGGATTCGCCGCCATCCGCACAGAgTTCAGAATGGCGTGTCCAGGTGAAGGTCTGCCTCCAGAGCTA
1051  F  L  L  S  I  I  R  Y  A  P  S  Q  E  W  V  E  I  H  L  S  Q  L  S  K  Q  M  A  D  G  F  A  A  I  R  T  E  F  R  M  A  C  P  G  E  G  L  P  P  E  L

3301 GTCCGAGACATGTTCCATAGCAGCAGGTGGACAAGCCCTGAAGGTTTAGGTCTAAGCGTATGTCGAAAGATTTTgAAGCTAATGAACGGTGAGGTTCAATACATCCGAGAATCAGAACGGTCCTATTTCCTCATCATTCTGGAACTCCCT
1101  V  R  D  M  F  H  S  S  R  W  T  S  P  E  G  L  G  L  S  V  C  R  K  I  L  K  L  M  N  G  E  V  Q  Y  I  R  E  S  E  R  S  Y  F  L  I  I  L  E  L  P

3451 GTACCTCGAAAGCGACCATTGTCAACTGCTAGTGGAAGTGGTGACATGATGCTGATGATGCCATAT
1151  V  P  R  K  R  P  L  S  T  A  S  G  S  G  D  M  M  L  M  M  P  Y
```

**Fig. S3 The eYHB DNA and protein sequences and residues (colored and bold) for potential nonsense mutation by EMS mutagenesis.** 25 synonymously ablated restriction sites within the eYHB coding sequence are underlined with the changed nucleotide being written in lowercase. The hairpin/tongue region of the PHY domain (W563 ~ A587) is also underlined. Since EMS elicits specific G/C-to-A/T mutation, five codons of Arginine (R), CGT, CGC, CGG, AGA and AGG, cannot be mutated into stop codons by one nucleotide change. These 52 non-targeted Arginine residues and their codons are marked with gray shade. The remaining 13 CGA-encoded Arginine, and 57 Glutamine (Q) as well as 11 Tryptophan (W) residues (total *n* = 81) are the targets for nonsense mutation. 26 residues identified as nonsense mutations in this study are boxed (*n* = 41, counting multiple independent mutation events on the same residues). Residues mutated multiple times are additionally indicated in italics and yellow highlighting.

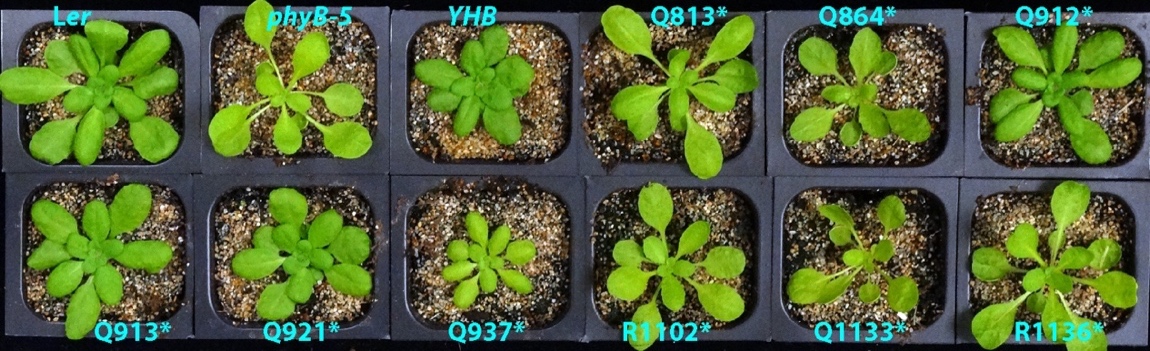

**Fig. S4 Adult plants of C-terminal nonsense mutations grown under short-day conditions for 36 days.** The Q912*, Q913*, Q921* and Q937* mutants (N-terminal to the HKRD domain) exhibit more compact rosette than the *phyB-5* mutant and are similar to L*er* WT, revealing residual activity of the truncated YHB in these four nonsense mutant lines.

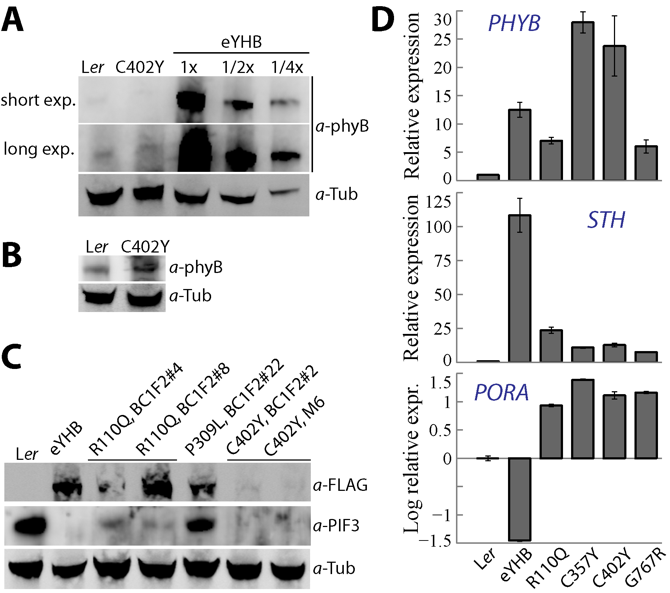

**Fig. S5 Reduction of steady state YHB protein levels by the C402Y mutation is likely due to protein instability, not by reduced transcription**. (**A**) Immunoblotting assay shows the phyB/YHB levels of both L*er* WT and the C402Y variant are at least eight-fold less than that of the eYHB parental line. (**B**) Another independent immunoblotting assay shows the YHB^C402Y^ level is comparable to the endogenous phyB level of L*er* WT. (**C**) After genetic background cleanup (backcross with *phyB-5*), the R110Q variant restores steady protein levels comparable with the eYHB parent and another variant P309L, whereas C402Y still retains drastically reduced protein level; homozygous BC1F2 lines of individual variants were tested; M6, the 6^th^ generation of propagated C402Y mutant. (**D**) Real-time PCR quantification of transcript levels of *PHYB*, *STH* and *PORA* genes; expression values are presented as mean ± S.D. (*n* = 3). Note that the C402Y and C357Y variants have higher *PHYB* transcript levels than the eYHB parent.

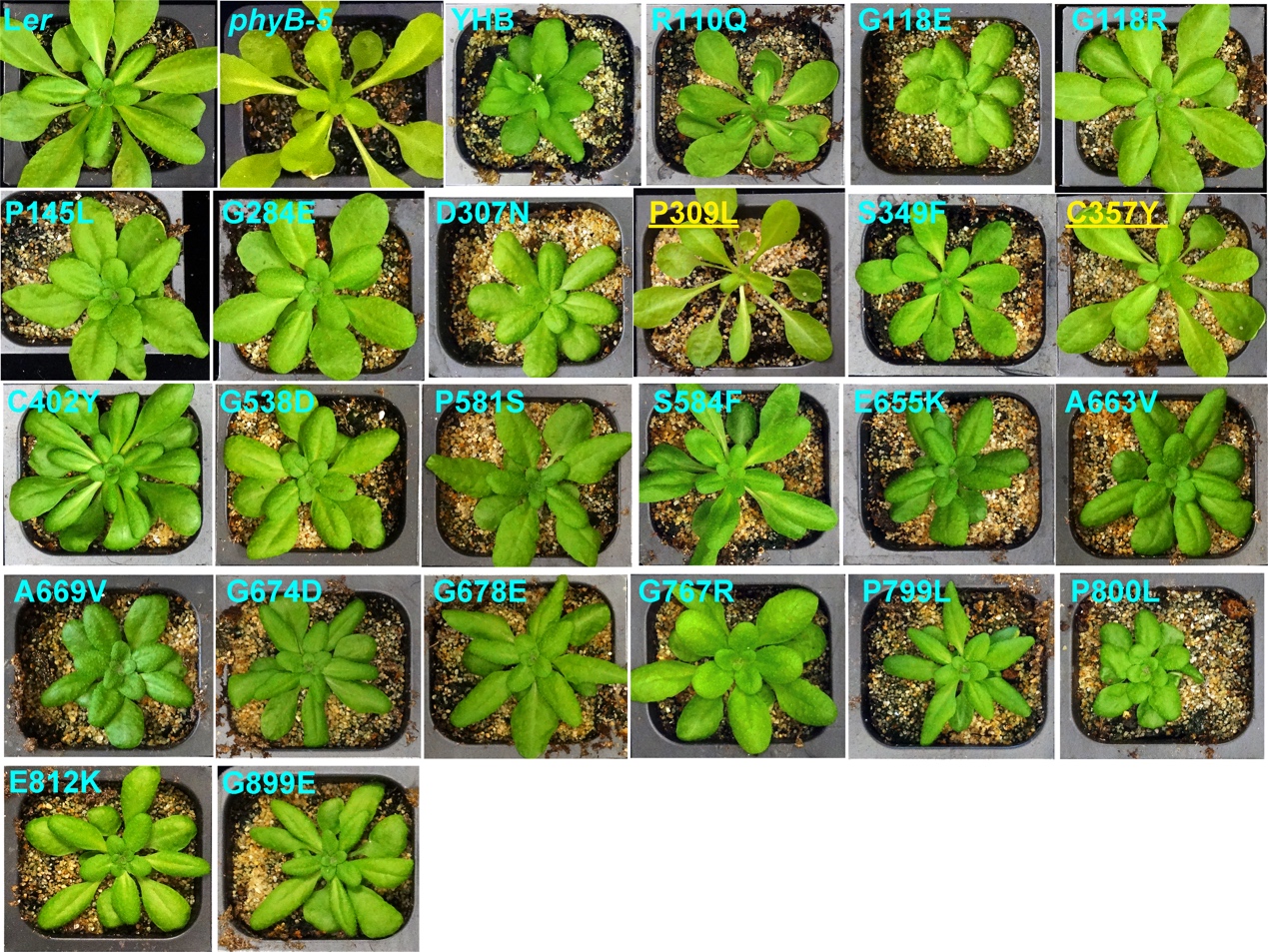

**Fig. S6 Missense suppressor plants grown under short-day conditions for 40 days.** Two suppressor lines restoring the long-petiole and small-bladed-leaf phenotype of the *phyB-5* mutant are indicated by underlining and yellow color.

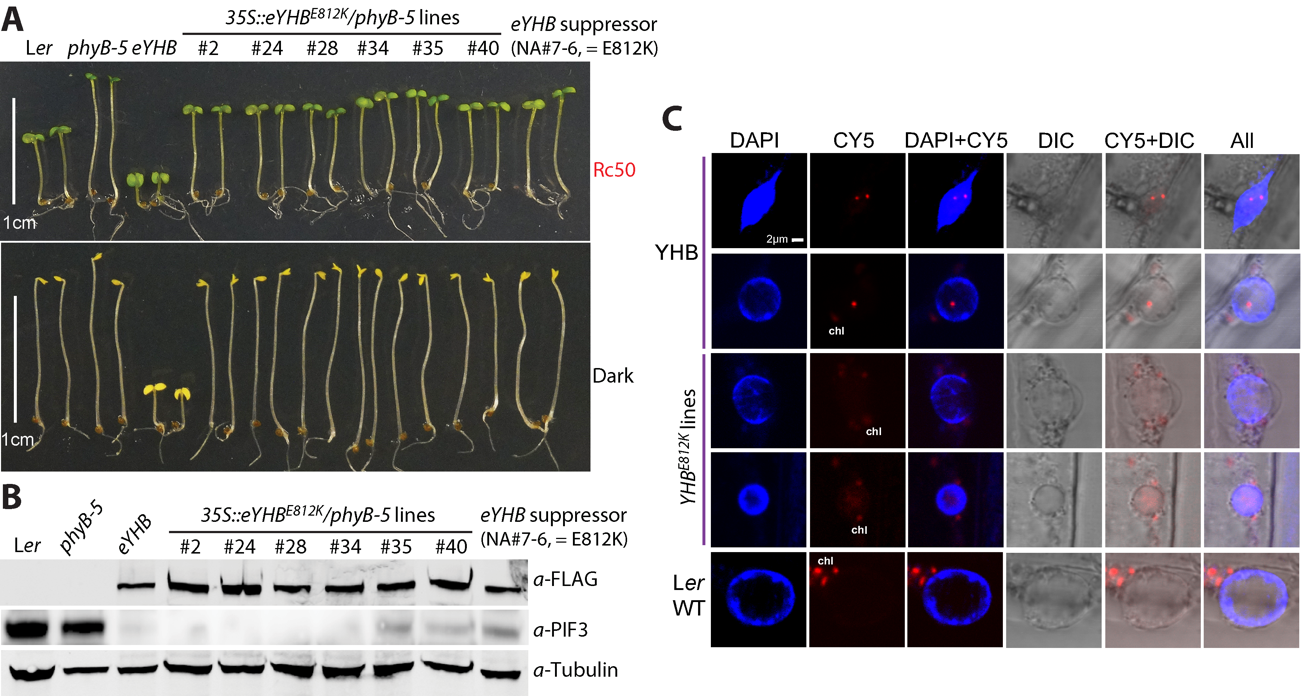

**Fig. S7 Validation of E812K suppressive effect on the YHB function by independent transgenic lines.** (**A**) Representative 4-day-old seedlings grown under continuous red light (50 µmol m^-2^ s^-1^) or in the dark. eYHB = eYHB-FLAG, six independent *35S::eYHB^E812K^/phyB-5* lines were tested. (**B**) Immunoblotting assay of dark-grown seedlings for the eYHB-FLAG (variant) and PIF3 protein levels; Note that due to membrane blotting issue, weak PIF3 bands were mistakenly missed from lines #2, #24, #28 and #34 compared to other YHB^E812K^ germplasms. (**C**) Confocal microscopy reveals a few large nuclear photobodies of YHB versus no nuclear fluorescence signal or faint diffuse nuclear distribution of *YHB^E812K^*.

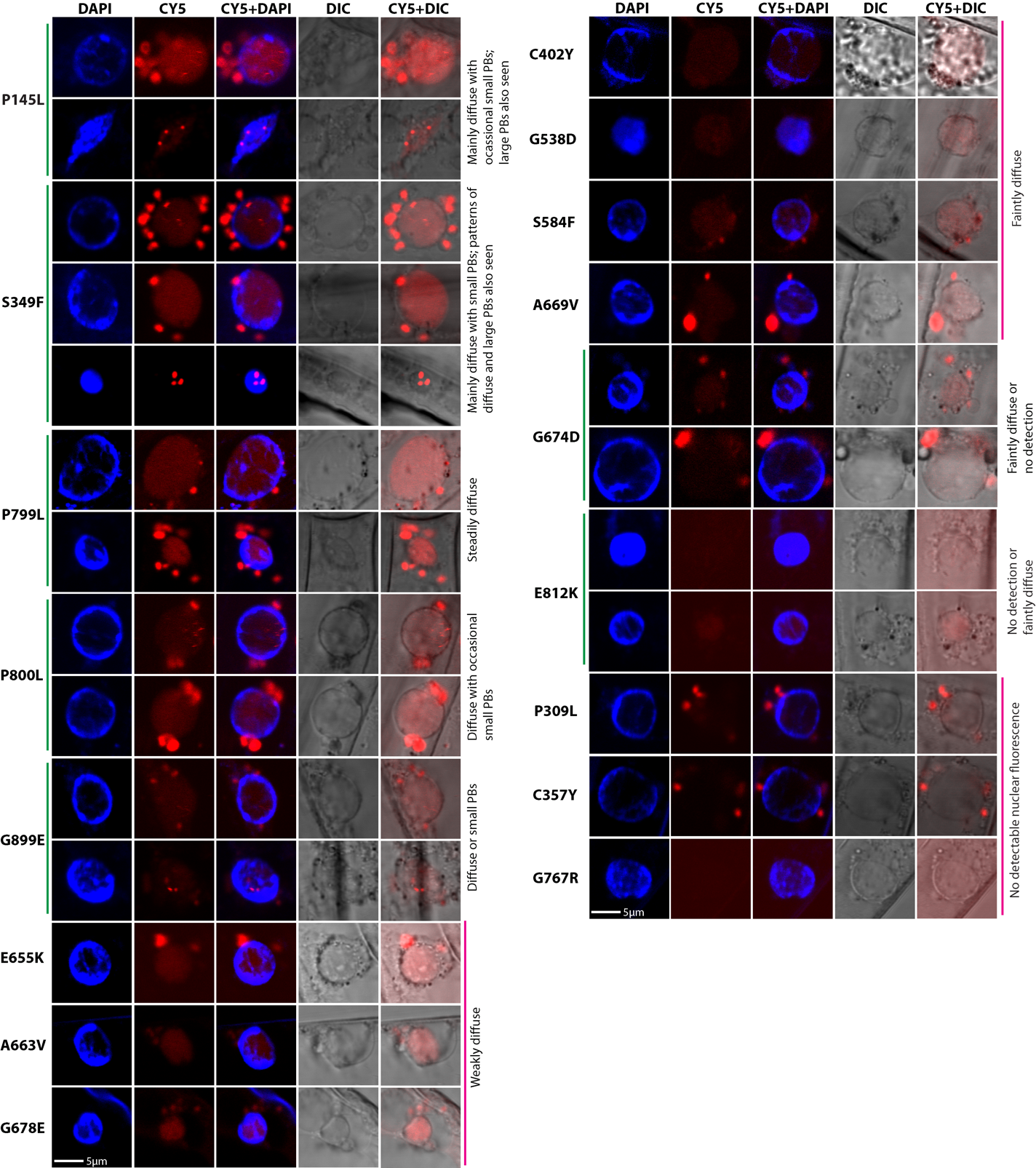

**Fig. S8 Confocal microscopy examination of subnuclear distributions of missense variants from dark-grown seedlings**. DAPI staining reveals the nucleus, the CY5 channel reveals red fluorescence emitted from YHB (if any) and chlorophyll as well. One mutant variant may have more than one distribution patterns, all of which are present. This figure is supplemental data for Fig. 4.

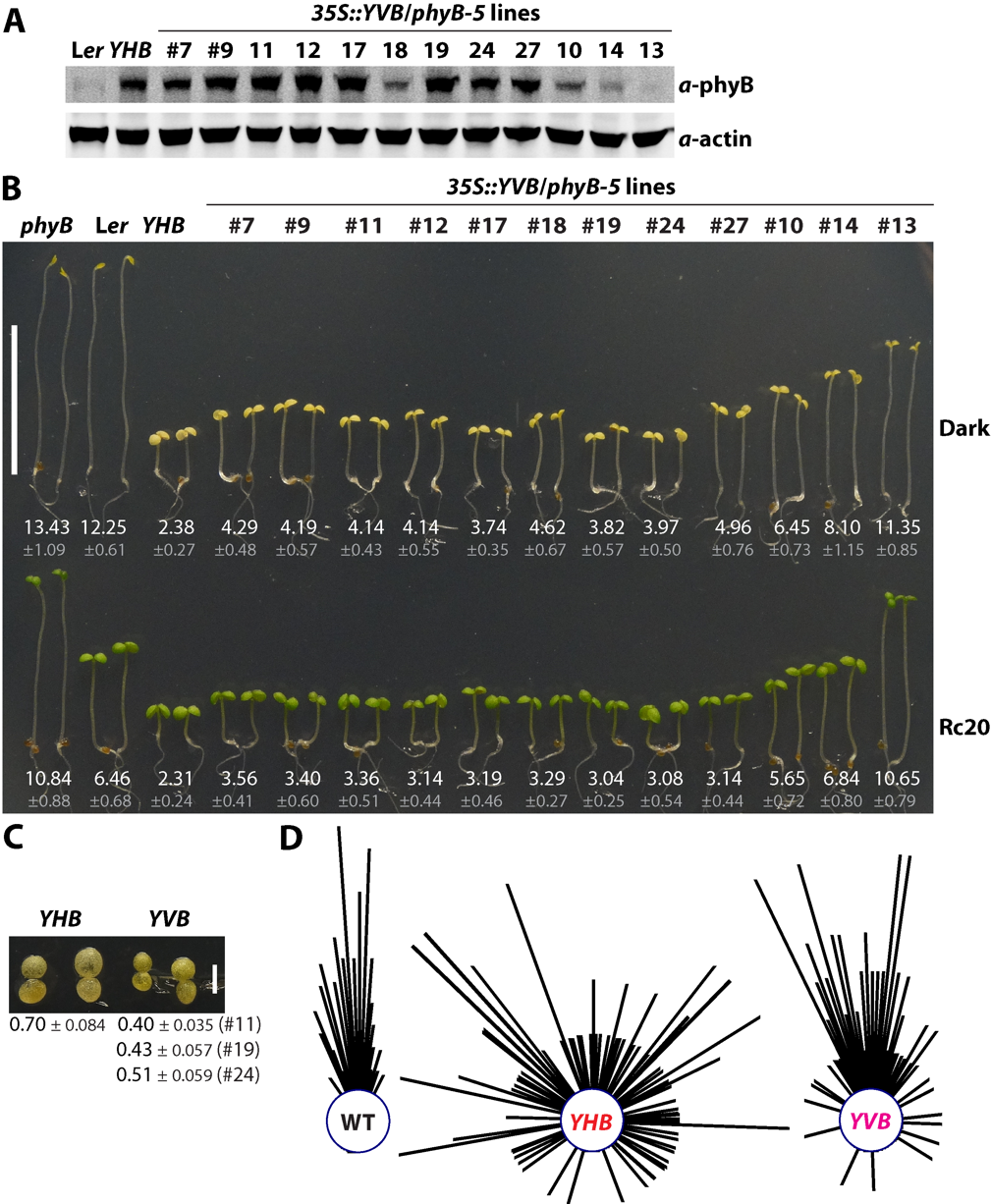

**Fig. S9** **The constitutive activity of phyB^Y303V^ (YVB) is weaker than phyB^Y276H^ (YHB**). (**A**) Immunoblot assay of YVB protein levels in dark-grown transgenic lines. (**B**) Five-day-old seedlings grown in the dark or continuous red light (20 µmol m^-2^ s^-1^), hypocotyl lengths are presented as mean ± S.D. (*n* = 20), scale bar = 1cm, note that lines #10, #14 and #13 with less YVB protein levels exhibit weaker cop phenotypes than other lines. (**C**) The cotyledon sizes of *YHB* seedlings are larger than those of *YVB* grown in the dark; cotyledon area (mm^2^) of 5-day-old seedlings are presented as mean ± S.D. (*n* = 20). (**D**) Circular histograms depict seedling growth orientation in the dark on vertically positioned plates, *n* = 100, examined *YVB* lines #9, #11, #19 and #24 exhibit similar orientation profiles, and line #9 is used for presentation.

**Supplementary Table S1. Identified mutation alleles in this study**

| Mutation Allele | Suppressor seedling phenotype | Sequenced mutant lines | Note and references |
| --- | --- | --- | --- |
| R107* | Null | NA#30-1, -2, -4, -6, -7 | New |
| R110* | Null | NB#64-1, -2 | New |
| W191* | Null | NA#3-5; NA#3-6-1; NA#4-1;  NB#54-1; NB#56-1; NB#57-1, -2, -3, -9, -15 | New |
| Q235* | Null | NB#17-1 | New |
| Q318* | Null | NA#35-12, -35, -36; | New |
| R322* | Null | NB#18-1, -2, -4, -5; NB#79-6 | New |
| W397* | Null | NB#22-1 | = *phyB-9*/*phyB-104* (Reed et al., 1993; Bradley et al., 1996) |
| Q423* | Null | NA#6-2; NA#47-1, -3, -18, -19, -20, -21, -26; NB#26-1 | New |
| Q438* | Null | NA#16-1, -2, -3; NB#2-1; | New |
| Q448* | Null | NA#32-1, -2, -3 | = *phyB-1* (Reed et al., 1993; Elich and Chory, 1997) |
| Q466* | Null | NB#138-2 | New |
| R554* | Null | NA#40-1, -2; | New |
| W563* | Nearly null | NA#14-1, -2; NA#36-24, -25; | New |
| Q577* | Null | NA#7-7 | New |
| W599* | Null | NA#18-1, -2; NB#1-1, -2; NB#41-1 | New |
| W679* | Nearly null | NB#45-1 | New |
| Q784* | Nearly null | NB#65-1 | New |
| W813* | Nearly null | NB#5-2; NB#65-2 | New |
| Q864* | Null | NB#25-1 | New |
| Q912* | Strong | NA#39-2, -5, -6, -7; | New |
| Q913* | Strong | NA#47-45, -47; | New |
| Q921* | Strong | NA#15-2, -3 | New |
| Q937* | Strong | NA#4-2, -3 | New |
| R1102* | Nearly null | NB#23-1 | New |
| Q1133* | Nearly null | NB#19-1, -4; NB#20; NB#33-1; NB#36-3 | New |
| R1136* | Nearly null | NB#12-1 | (Wagner and Quail, 1995) |
| R11OQ | Moderate ~ strong | NA#17-1 | (Oka et al., 2008; Kikis et al., 2009) |
| G118E | Moderate | NA#4-4, -6 | New |
| G118R | Strong | NB#9-1, -2, -5; NB#29-3, -11 | (Krall and Reed, 2000; Chen et al., 2003; Oka et al., 2008). |
| P145L | Weak ~ moderate | NA#14-4, -6, -7 | New |
| G284E | Null in darkness, weak ~ moderate in the light | NB#16-1, -2; NB#123-1 | (Oka et al., 2008) |
| D307N | Weak ~ moderate | NA#43-1, -2 | New; strong mutation D307A reported (Zhang et al., 2013) |
| P309L | Null | NB#62-1 | (Oka et al., 2008) |
| S349F | Strong | NA#50-1, -2; | = *phyB-102* (Bradley et al., 1996) |
| C357Y | Nearly null | NB#21-1, -2, -3; NB#22-9; NB#31-2; NB#34-10 | Chromophoreless. New. C357A/S reported (Wagner et al., 1996; Kircher et al., 1999; Oka et al., 2011) |
| C402Y | Nearly null | NB#27-7 | New |
| G538D | Strong | NA#5-2, -3; NA#37-1, -2; | New |
| P581S | Moderate ~ strong | NB#5-1, -5, -6 | New |
| S584F | Nearly null | NB#94-1, -2 | (Oka et al., 2008) |
| E655K | Moderate | NA#30-16, -17; | New |
| A663V | Moderate | NA#32-13 | New |
| A669V | Moderate | NA#16-13  NB#128-1 | New |
| G674D | Moderate ~ strong | NA#6-4; NA#8-6, -7, -10, -11; NA#36-38; | (Chen et al., 2003) |
| G678E | Moderate ~ strong | NA#45-1; NB#8-1, -2, -4; NB#17-2, -3; NB#22-5; NB#23-2; NB#65-5 | New |
| G767R | Strong | NB#21-17; NB#22-11; NB#34-3 | (Wagner and Quail, 1995; Matsushita et al., 2003; Hu and Lagarias, 2024) |
| P799L | Moderate | NA#31-3; NB#103-1 | New |
| P800L | Weak ~ moderate | NA#15-5 | New |
| E812K | Nearly null | NA#7-6; NB#104-5, -6; NB#107-1, -2, -3, -4; NB#109-1, -2 | = *phyB-101* (Wagner and Quail, 1995; Bradley et al., 1996; Elich and Chory, 1997; Chen et al., 2003) |
| G899E | Strong | NB#1-4, -5; NB#19-5, -6; NB#61-3 | New |

**Supplementary Table S2. PHYB missense mutations identified or characterized from previous and current work** (in a separate excel file)

**Supplementary Table S3. Primers used in the current study**

| Primer ID | Primer sequence (5’ – 3’) | Purpose |
| --- | --- | --- |
| oWH202 | CCGACAGTGGTCCCAAAGATGGA | YHB-3xFLAG transgene amplification |
| oWH307 | CTGGTGTGTGCGCAATGAAA | YHB-3xFLAG transgene amplification |
| PHYB-R3 | CCATAGCTTCTTCAACTGAGAG | Used with oWH202 for amplifying partial transgene fragment |
| oWH295 | at*ggtacc*ATGGTTTCCGGAGTCGGG | PHYB N651 fragment amplification |
| oWH314 | agt*cccggg*TGCACCTAACTCATCAATCCC | PHYB N651 fragment amplification |
| oWH330 | GTGCTGTTCAATCGCAGAAA | *PHYB* qRT-PCR |
| oWH331 | TCCACGACAGTGTCACACAA | *PHYB* qRT-PCR |
| oWH61 | CAGAGTCTCTCTAAACCGCCAACT | *STH* qRT-PCR |
| oWH62 | TCATCGGTAGCCCACAAAGG | *STH* qRT-PCR |
| oWH242 | CCAACGGCTAATCAACCTACTTTC | *PORA* qRT-PCR |
| oWH243 | AATGGTTTATCCCAACGCTAAGC | *PORA* qRT-PCR |
| oWH45 | GGCCTTGTATAATCCCTGATGAA | *UBQ10* qRT-PCR |
| oWH46 | AGAAGTTCGACTTGTCATTAGAAAGAAA | *UBQ10* qRT-PCR |
